# The landscape of transcriptional and translational changes over 22 years of bacterial adaptation

**DOI:** 10.1101/2021.01.12.426406

**Authors:** John S. Favate, Shun Liang, Srujana S. Yadavalli, Premal Shah

**Affiliations:** Department of Genetics, Rutgers University, Piscataway NJ, USA; Human Genetics Institute of New Jersey, Rutgers University, Piscataway NJ, USA; Waksman Institute, Rutgers University, Piscataway NJ USA

## Abstract

Organisms can adapt to an environment by taking multiple mutational paths. This redundancy at the genetic level, where many mutations have similar phenotypic and fitness effects, can make untangling the molecular mechanisms of complex adaptations difficult. Here we use the *E. coli* long-term evolution experiment (LTEE) as a model to address this challenge. To bridge the gap between disparate genomic changes and parallel fitness gains, we characterize the landscape of transcriptional and translational changes across 11 replicate populations evolving in parallel for 50,000 generations. By quantifying absolute changes in mRNA abundances, we show that not only do all evolved lines have more mRNAs but that this increase in mRNA abundance scales with cell size. We also find that despite few shared mutations at the genetic level, clones from replicate populations in the LTEE are remarkably similar to each other in their gene expression patterns at both the transcriptional and translational levels. Furthermore, we show that the bulk of the expression changes are due to changes at the transcriptional level with very few translational changes. Finally, we show how mutations in transcriptional regulators lead to consistent and parallel changes in the expression levels of downstream genes, thereby linking genomic changes to parallel fitness gains in the LTEE. These results deepen our understanding of the molecular mechanisms underlying complex adaptations and provide insights into the repeatability of evolution.

## Introduction

Comparative genomic approaches and large scale mutation experiments have allowed us to map genetic changes to phenotypic changes underlying adaptation in many cases involving individual genes such as hemoglobin^1^, hormone receptors^2^, and influenza proteins^3,4^. However, when organisms adapt to novel environments such as during yeast evolution under nutrient limitation^5–7^, adaptation to high-temperature stress^8^, bacterial evolution during infections^9^, and long-term adaptation of *Escherichia coli* to minimal media^10–13^, genomic changes are widespread. Understanding how these changes lead to functional changes at the molecular level is critical to understand the mechanistic basis of adaptations.

Here we use the *E. coli* long-term evolution experiment as a model system to characterize the mechanistic basis of adaptation to a novel environment. Recent studies using LTEE have quantified the dynamics of fitness growth^14^, identified the proportion of beneficial mutations^12^, characterized mutational dynamics in the system^13^, and identified the mechanistic basis of specific adaptations such as citrate utilization in Ara-3^15^. Despite significant contributions to the understanding of adaptation in the LTEE, the role that changes in transcription and translation play in increasing growth rates remains unexplored. An earlier study of gene expression changes in LTEE showed parallel changes in transcription profiles in two of the twelve evolved lines, Ara-1 and Ara+1, at 20,000 generations using radioactive microarrays^16^. Whether parallelism in gene expression changes extends to the other lines and persists over a more extended period remains unknown. Furthermore, since changes at the transcriptional level can be buffered at the translational level^17,18^, changes to both must be considered. Finally, significant changes in cell-size and morphology of the bacteria^19,20^ over the course of adaptation indicate a need to quantify both relative and absolute changes in expression.

## Results

To address these questions, we performed RNA-seq and ribosomal footprinting (also called Ribo-seq)^21^ in the exponential phase of the ancestral strains and single clones from each of the 12 evolved lines at 50,000 generations (Fig. 1A). We analyzed single clones from Tenaillon et al. 2016 and considered 4216 protein-coding genes from the ancestors. We aligned sequencing data for each evolved clone to its unique genome. We restricted our analysis to 11 out of 12 evolved lines due to ancestral contamination in one of our samples (Ara+6). We averaged between 151 and 1693 reads per gene across the 52 libraries (Fig. S1A, Table S1). The distributions of read counts per gene were similar across lines, replicates, and sequencing methods (Fig S1C). We also observed a clear three-nucleotide periodicity in our Ribo-seq datasets (Fig. S1B, Table S2).

**Figure 1:**
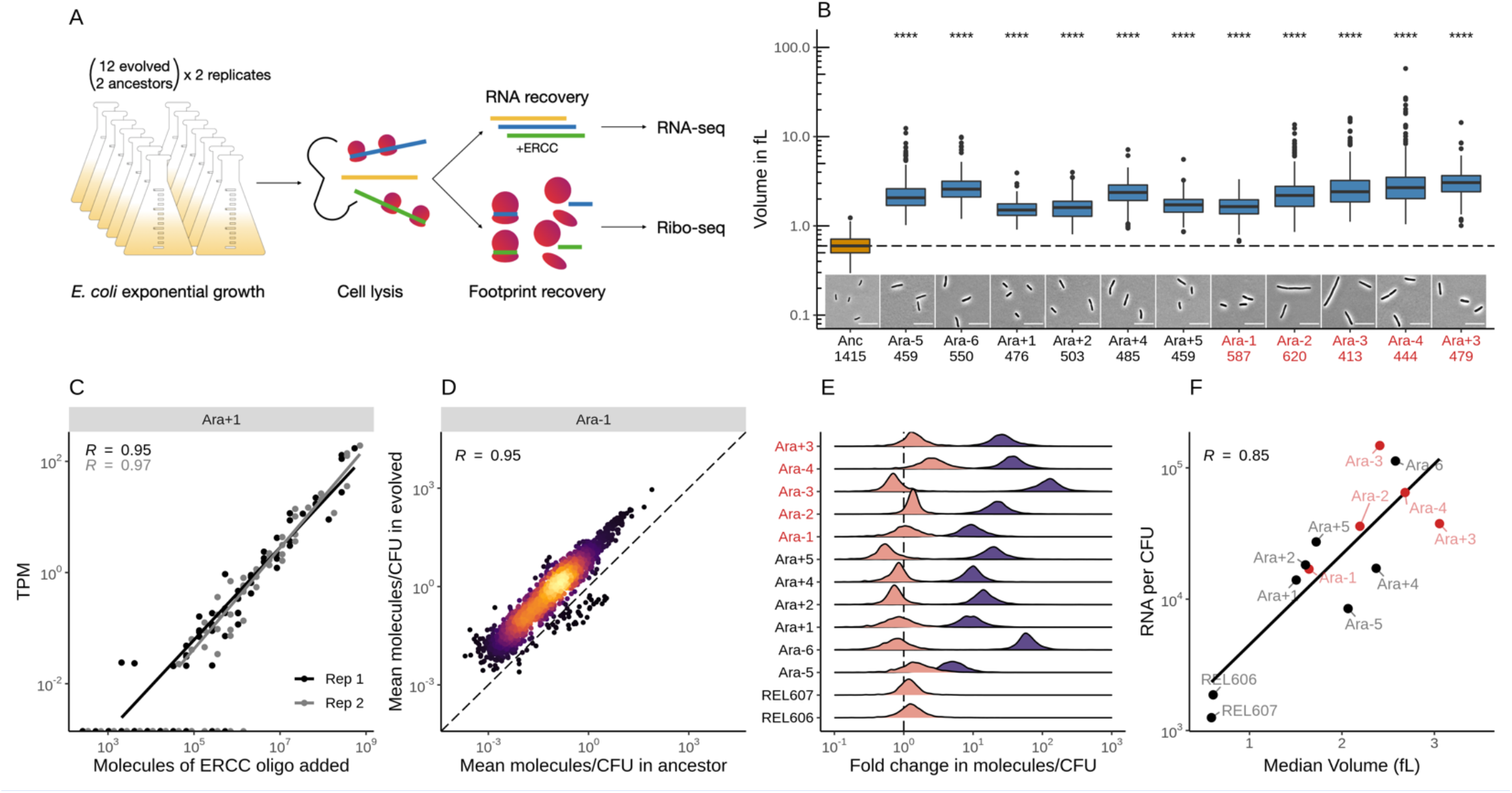
**A**. Schematic of the experimental design. **B**. All evolved lines are larger than the ancestral strain. Distributions of cellular volume as determined by phase-contrast microscopy and assuming sphero-cylindrical shape of *E. coli* along with representative images for each line. Numbers underneath a line’s name indicates the total number of cells imaged (scale bar is 10um, see Figure S3 for representative images.). The dashed line indicates the ancestral median, p-values indicate the results of a t-test when each line is compared to the ancestor, **** p ≤ 0.0001. Lines listed in red have mutator phenotypes. **C**. Spike-in RNA control abundances are correlated with their estimates in sequencing data. Linear models relating the number of molecules of each ERCC control sequence added to their RNA-seq TPM (transcripts per million) in Ara+1 RNA-seq samples (see Fig. S4 for all lines). **D**. Most genes have a higher absolute expression in evolved lines. Changes in the absolute number of mRNA molecules per CFU (colony forming unit) in the 50,000th generation of Ara+1 relative to the ancestor. The values plotted are the average between 2 replicates of the evolved lines and both replicates from both ancestors (REL606 and REL607; see Fig. S4 for all lines). **E**. Absolute changes in mRNA abundances in evolved lines are significantly larger than the variation between biological replicates (t-test, p < .0001 in all cases). Distributions of fold-changes of mRNA molecules per CFU. Pink curves indicate gene-specific fold-changes between biological replicates for each line (centered around 1). Purple curves show the fold-change from the 50,000th generation of an evolved line to the ancestor. Fold-change was calculated in the same manner as in D. **F**. Larger evolved lines have more mRNA per CFU. Relationship between the median volume for each line and the total number of RNA molecules per CFU for each line. Total molecules of RNA are calculated as the sum of the average number of molecules for each gene between replicates.

### Evolved lines are larger and carry more mRNAs

Contrary to expectations, every evolved line in the LTEE has become larger in size compared to the ancestor^19,20,22^. While bacterial size (cell volume) is a function of its growth rate, which typically depends on nutrient availability^23–25^, the increase in cell size in LTEE is not entirely a consequence of faster growth rate^19^. This increase appears to be under selection and is partly caused by mutations in Penicillin-binding protein genes, which also led to the increased circularity of the cells^26^. Moreover, cultures of the evolved lines were recently found to have higher biomass with proportionally higher amounts of nucleic acids compared to the ancestors^27^. Because changes to cell volume can affect transcription rates and alter relative concentrations of RNA molecules^28^, we chose to quantify changes in the absolute abundance of mRNAs.

We used phase-contrast microscopy to measure the size and shape of cells in each of the ancestral and evolved lines and calculated cell volume based on these measurements (see methods, Table S3). We find that each evolved line has a larger volume than the ancestor (Welch’s t-test, p < 0.0001 for all lines) (Fig. 1B). We also find that evolved lines form filaments more frequently and formed longer filaments than the ancestor (see Supplementary Analysis). However, the larger size of evolved lineages is not entirely due to higher filamentation. Even after filtering out filaments (cells >3x median volume), all evolved lines were still significantly larger compared to the ancestor (Welch’s t-test, p < 0.0001 for all lines) (Fig. S2B).

To measure how changes in cell size affect absolute RNA abundances, we measured the number of colony-forming units (CFU) that went into each library (Table S4). We added the ERCC RNA spike-in controls^29^, a set of 92 RNA oligos in known amounts, to our RNA-seq libraries (table S5). This allowed us to quantify the number of molecules per CFU for each transcript. We find a linear relationship between the number of molecules of ERCC oligos and the number of transcripts quantified using RNA-seq (TPM) (Fig. 1C, S4A). Fold-changes in absolute counts ranged widely in each of the lines (Fig. 1E, Table S6) but were overwhelmingly greater than one. Moreover, the increase in mRNA abundances in evolved lines relative to the ancestor were greater than differences in abundances between corresponding biological replicates (Fig. 1E, t-test, p < .0001 in all cases). This suggests that all evolved lines have more mRNA molecules compared to the ancestral strains. Finally, we show that evolved lineages with larger cells have more mRNAs (Fig. 1F), suggesting that absolute abundances of mRNAs scale with cell size.

### Gene expression changes are parallel at both transcriptional and translational levels

Despite a high degree of parallelism in fitness, few mutations are shared across the evolved lineages, and each of the lines was founded on a unique set of mutations^12^. At the gene level, only 57 genes have mutations in two or more lines^12^. Moreover, it remains unclear if the functional effects of these mutations are similar across lines. To bridge the weak parallelism at the genotypic level with the strong parallelism at the fitness level, we took gene expression as a molecular phenotype and quantified transcription and translation. Earlier radioactive microarray-based experiments with two evolved lineages (Ara+1 and Ara-1) at 20,000 generations have showed that the expression patterns between the two evolved lines were more similar to each other than either were to the ancestor^16^. However, it remains unclear if the pattern of parallel gene expression changes is identical across all evolved lineages and has remained mostly parallel over a more extended period.

We find that expression levels of genes were surprisingly similar across evolved lineages. Pairwise correlations based on TPM showed a high degree of similarity among the evolved lines for RNA-seq and Ribo-seq datasets (Fig. S5A, Table S1). Interestingly, pairwise correlations between evolved lines were not significantly different from correlations between evolved lines and the ancestors (Fig. S5B). This suggests that expression patterns of many genes remained mostly unchanged over 50,000 generations. We then sought to systematically quantify the degree of expression changes in both RNA-seq and Ribo-seq datasets using DESeq2^30^ in each of the evolved lines (Table S7). Overall, half of all genes across all lines had less than a 30% change in their expression levels (Fig. S5C). However, several genes showed large changes in their expression patterns that varied by a thousand-fold (log2 fold-change > 10).

We find a high degree of parallelism in expression changes at both the transcriptional and translational levels (Fig. 2 and S6). The top 100 up and downregulated genes (defined as having the largest mean positive or negative fold-change across the evolved lines) showed remarkably similar fold-changes (RNA-seq, Fig. 2A; Ribo-seq, Fig. S6A). Distributions of all pairwise comparisons of fold-changes in evolved lines showed positive correlations, which became even more positive when considering only statistically significant genes (RNA-seq, Fig. 2B; Ribo-seq, Fig. S6D). Interestingly, we find that a higher number of genes were downregulated than upregulated across most lines (RNA-seq, Fig. 2E; Ribo-seq, Fig S6C). Moreover, the magnitude of downregulations was larger than that of upregulations in all but Ara+3 (Welch’s t-test, p < 0.05 in all cases) (RNA-seq, Fig. 2B; Ribo-seq, Fig S6B). Surprisingly, evolved lines arrived at similar transcriptional and translational profiles regardless of whether they had a mutator phenotype or not (Fig. S6E).

**Figure 2:**
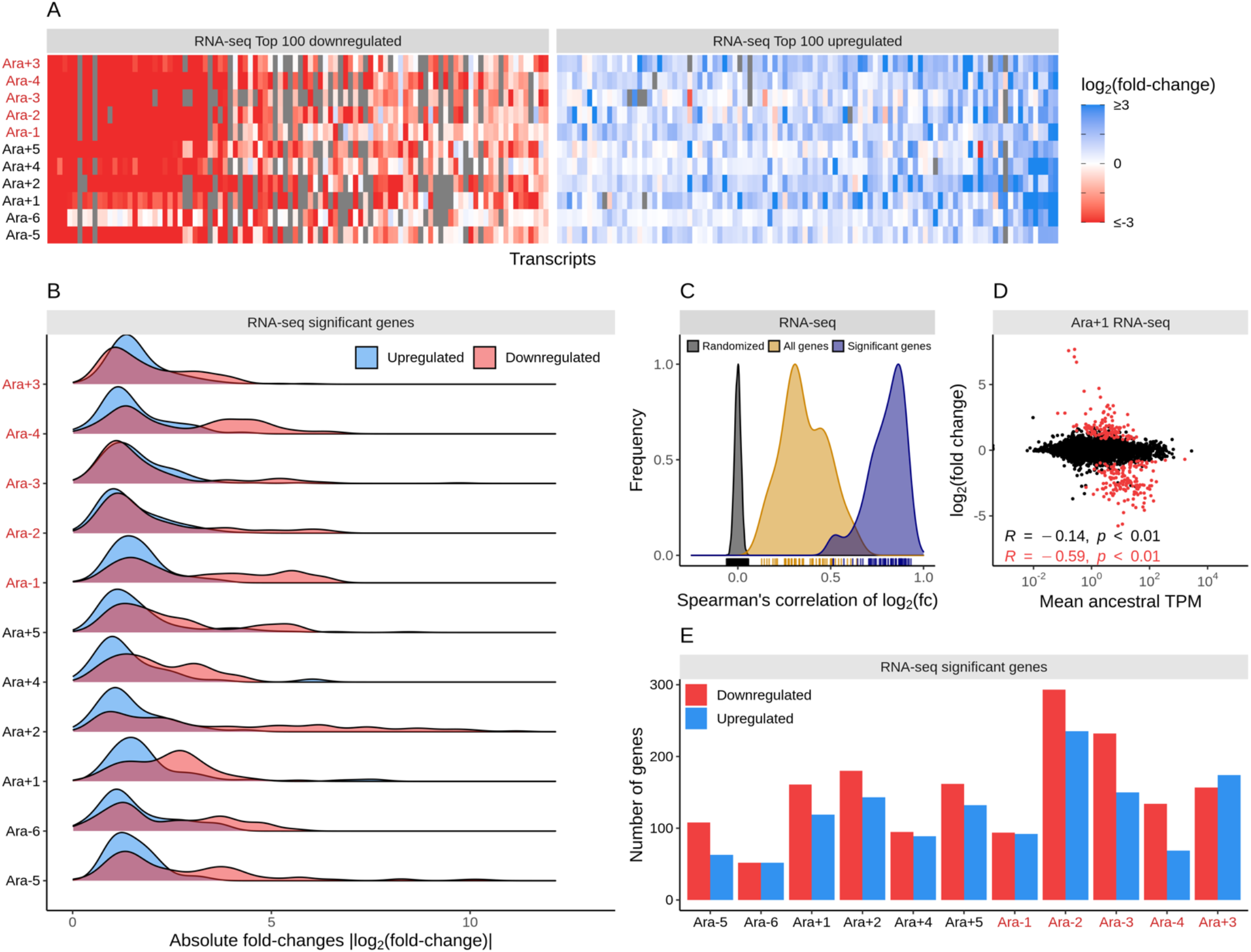
**A**. Parallelism in expression changes across evolved lines. The fold-changes of top 100 down and upregulated genes in each of the lines in the RNA-seq datasets. Genes are ordered from left to right in order of increasing mean fold-change across evolved lines. Gray bars represent gene deletions. **B**. Downregulated genes have larger effect sizes than upregulated genes. Distribution of statistically significant fold-changes in each line. Statistical significance was based on DESeq2 results using q ≤ 0.01. **C**. Pairwise correlations of evolved lines based on all (yellow curve) or only statistically significant (blue curve) RNA-seq fold-changes. Each of these curves is significantly different from a distribution based on correlations made after randomizing the fold-changes (grey curve) within each line (p ≤ 0.01, t-test). **D**. Fold-changes in expression levels of genes in evolved lines scale negatively with their ancestral expression levels. The relationship between ancestral TPM in the RNA-seq dataset and RNA-seq fold-change in Ara+1. The red dots represent significantly altered genes, and the black dots represent the remaining genes. **E**. The number of significantly down and upregulated genes in each line.

We next examined if changes in expression levels of a gene were somehow related to their expression in the ancestor. When we considered all genes, we observed a weak negative relationship between ancestral TPM and fold-change in an evolved line (Fig. 2D, S6F). This negative relationship is likely a by-product of the overall increase in mRNA abundances with cell-size. Due to biophysical constraints, genes with high ancestral expression are unlikely to see large increases in mRNA abundances relative to genes with low expression. As a result, genes with low ancestral mRNA abundances appear more upregulated when considering only relative expression levels. However, when only statistically significant genes were considered, we see a very strong negative relationship in most lines. The slope of this relationship is distinctly more negative than for all genes. Additionally, the proportions of significantly upregulated genes decreased with the ancestral gene expression level for most lines (Fig. S6G).

We observed high levels of parallelism in expression changes despite few shared mutations across multiple lines (Fig. 3A, Table S8). We find that both the proportions of downregulated genes and their magnitude of downregulation increased with the number of lines a gene was significantly altered in (Fig. 3C and D), indicating that more downregulations were shared across lines than upregulations. This implies that there are fewer genes and pathways whose downregulation increases fitness, whereas genes and pathways whose expression increases enable higher fitness are more varied and unique to each line. We find similar patterns for the Ribo-seq datasets (Fig. S7).

**Figure 3:**
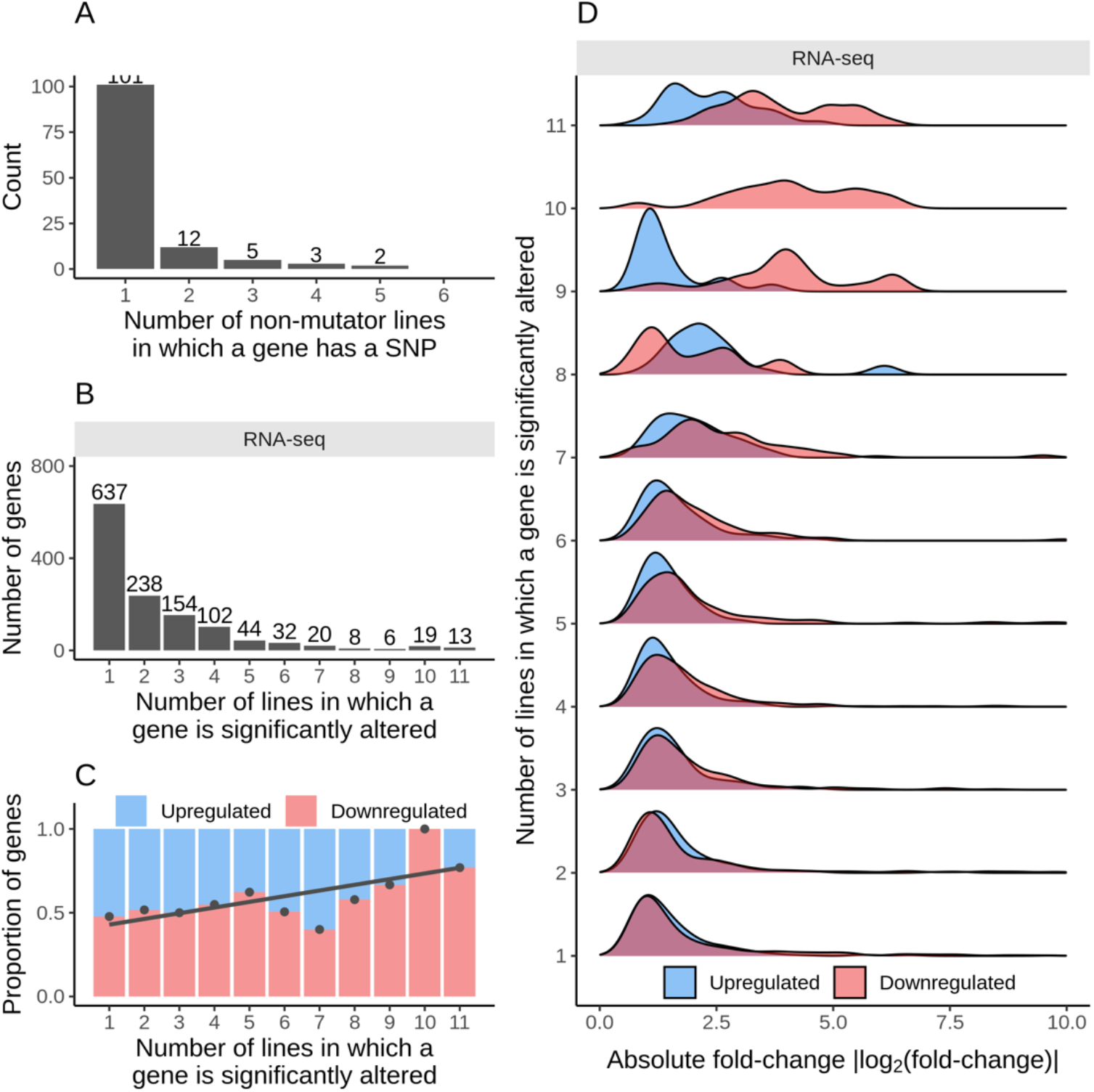
**A**. The number of non-mutator lines in which a gene has at least one SNP inside the coding sequence. **B**. The number of evolved lines in which a gene’s expression level was significantly altered (q ≤ 0.01) was based on the DESeq2 results for RNA-seq datasets. **C**. Frequently altered genes are typically downregulated. The proportion of up and downregulated genes as a function of their frequency of expression changes across lines. **D**. Frequently downregulated genes have larger effect sizes than upregulated genes. Distributions of the RNA-seq fold-changes for the genes in the x-axis categories of C.

### Transcriptional changes drive translational changes

Translational regulation affects the rate at which an mRNA produces its protein product. Different mRNAs are translated with varying efficiencies in both eukaryotes and prokaryotes^21,31,32^. However, the role of changes in translational regulation during adaptation and speciation remains poorly understood and is heavily debated^18,33^. To study translational changes, we performed high-throughput ribosome-footprinting in both the evolved lines and their ancestors.

Interestingly, we find that gene-specific ribosome-footprint abundances were highly correlated with mRNA abundances (R ≥ 0.92 for all lines, Fig. 4A and S8A). Since the number of ribosome-footprints from a gene also depends on its mRNA abundances, we used Riborex^34^ to evaluate gene-specific changes in ribosomal-densities in each of the evolved lines relative to the ancestor. Surprisingly, we find very little evidence of translational changes (Fig. 4B, Table S9). The number of genes with significantly altered (q ≤ 0.01) ribosome-densities ranged from 0-6 genes across all lines, with a total of only 18 unique genes showing altered ribosome-densities. Overall, changes in ribosome-densities on transcripts were sparse, suggesting that transcriptional changes are the dominant force behind expression changes in the LTEE.

**Figure 4:**
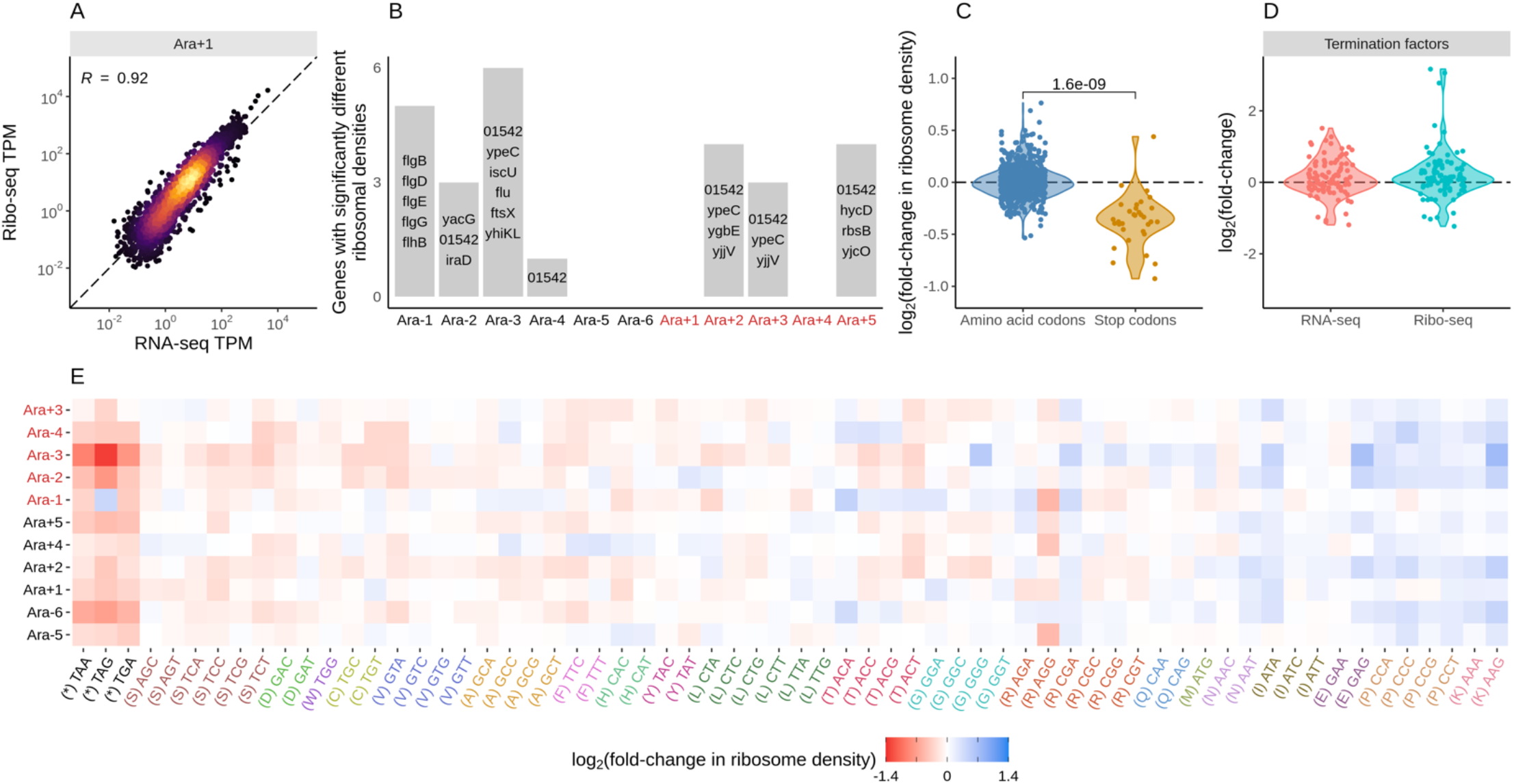
**A**. Translational changes are positively correlated with transcriptional changes. The relationship between RNA-seq and Ribo-seq TPM in Ara+1. The TPMs are averaged between the replicates. **B**. The distribution and identity of genes with significantly altered ribosomal densities (q ≤ 0.01). **C**. Evolved lines have faster translation termination. Stop codons had lowered ribosome density compared to amino acid codons. Each point represents a stop codon from an evolved line, and the y-axis is fold-change in ribosomal density relative to the ancestor. P-value based on a t-test. **D**. Fold-changes in expression levels of translation termination factors and related genes *ykfJ, prfH, prfA, prmC, prfB, fusA, efp, prfC*. **E**. Changes in codon-specific ribosome densities in each of the evolved lines relative to the ancestor. Codons are arranged from left to right in order of increasing mean fold-change for their respective amino acid across the lines.

While ribosome-density changes reflect changes to the overall number of ribosomes per transcript, they do not reveal information about the translation of specific codons or amino acids. We find that the ribosome-densities at stop codons were significantly lower in all the evolved lines than in the ancestors (Fig. 4C and 4E, Table S10), suggesting that translation termination was significantly faster in evolved lines. Translation initiation and termination are relatively slow processes compared to elongation. As a result, faster termination might be adaptive in that it allows faster recycling of ribosomes, thereby increasing overall protein synthesis rates. Furthermore, we reasoned that this change in stop-codon ribosome-densities might be due to changes in expression levels of proteins that aid translation termination, such as release factors. We examined changes in genes related to termination, namely *frr* (ribosome recycling factor^35^), *fusA* (elongation factor G^36^), *prfABC* (peptide release factors A, B, C^37,38^), and *prmC* (a methylase required for the function of prfAB^39^). These genes showed differing directions and magnitudes of alteration at the RNA level, and these changes were rarely statistically significant (Fig. 4D, S8C). *prfB* and *prfC* facilitate the release of a protein from the ribosome at a stop codon and were typically upregulated, indicating an increase in their expression might be responsible for faster translation termination.

We also find higher ribosome-densities at Proline codons across all lines, indicating that elongation rates at these codons have slowed. Given this apparent slowdown at proline codons, we examined if genes involved in proline biosynthesis had altered expression levels. However, the three enzymes directly involved in proline biosynthesis - *proA*, *proB*, and *proC*, the proline tRNA ligase - *proS*, and elongation factor P involved in alleviating ribosome pausing at polyproline motif ^40^, were not significantly altered in any of the lines (Fig. S8B). We suspect that the higher ribosome-densities at Proline codons are likely due to lower levels of charged proline tRNAs.

### Transcriptional and translational changes of frequently deleted genes

Large deletions are among the most frequent class of mutations in the LTEE^11,12^ and several gene deletions are shared across multiple evolved lineages (Fig. 5A). For example, the *rbs* operon is partially or entirely deleted in every evolved line, making them unable to catabolize ribose. This loss of *rbs* operon leads to increased fitness relative to the ancestor^41^. We also find that genes deleted entirely in at least four lines had lower expression in the ancestor (Fig. 5B). While the fitness benefit of specific deletions such as *rbs* operon has been experimentally validated, it is more challenging to systematically assess the effects of deletions in only some of the lines. This is especially true of the large deletions that encompass multiple genes of unrelated functions. Since downregulation and deletions of genes have similar functional effects (that is, removal of the gene product), we hypothesized that frequently deleted genes would be typically downregulated in lines where the gene was still present. Surprisingly, we find no enrichment in the downregulation of genes deleted in at least four lines (Fig. 5C). One reason for this lack of enrichment might be the mechanism by which most genes are deleted in LTEE. Deletions in LTEE are typically mediated by insertion-elements, spanning multiple kilobases and encompassing multiple genes (Fig. 5D). On average, 17 genes were lost per deletion event. Our results suggest that while deletions of a few genes within these large deletions might be under selection, most of other deletions are simply genetic hitchhikers.

**Figure 5:**
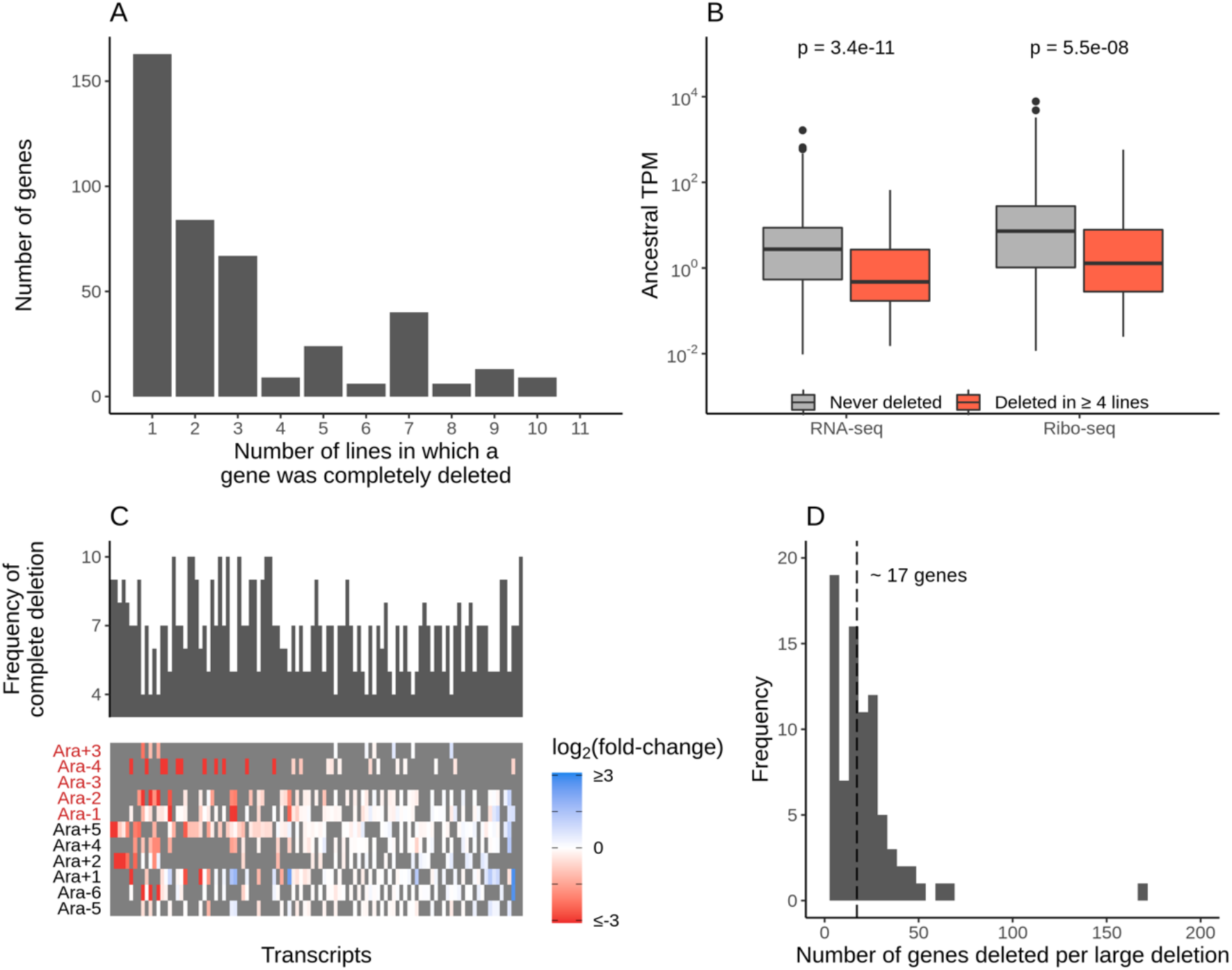
**A**. The frequency with which a gene was deleted entirely across the lines. **B**. Frequently deleted genes have lower expression levels in the ancestors. The distributions of ancestral TPMs of genes were deleted entirely in at least four lines (red) or were never deleted in any of the lines (grey). P-values based on a t-test. **C**. Frequently deleted genes are not typically downregulated in lines where they are present. Heatmap represents RNA-seq fold-changes of all genes deleted in at least four lines. Genes are ordered from left to right in order of increasing mean fold-change across evolved lines. Gray bars represent gene deletions. The histogram above the heatmap indicates the frequency of deletion of corresponding genes in the heatmap. **D**. Number of genes deleted per large deletion in LTEE across all 12 lines. The dashed line indicates the average number of genes deleted per deletion (~17).

### Functional characterization of differentially expressed genes

To identify functional categories and pathways that are altered as a result of expression changes in each line, we looked for enrichment in KEGG pathways^42^, gene ontology terms^43^, and pathway perturbation scores (PPS) from the BioCyc collection of databases^44^ (Fig. S10, Table S13, see methods for details on each). For these analyses, we considered deleted and pseudogenized genes as being downregulated.

Though many categories were altered across the lines in the KEGG analysis (see Table S11 for complete results), we chose to focus on those that were significantly altered (FDR ≤ 0.05) in at least four lines. We find a high degree of parallelism between the evolved lines for KEGG pathways that are significantly altered based on RNA-seq datasets (Fig. 6A, C; see Fig. S9A for Ribo-seq scores). Consistent with earlier microarray experiments, we find that the flagellar assembly genes are significantly downregulated^16^ in 10 out of 11 evolved lines. In addition, because the evolved lines are growing in a stable environment over evolutionary timescales, it stands to reason that genes involved in responding to stress and environmental changes will be downregulated. As expected, we find that genes associated with biofilm formation, two-component signaling pathways, and ABC transporters are all downregulated across most lines. Furthermore, we find that selection for faster growth in LTEE has led to significant increases in expression levels of genes involved in amino acid biosynthesis and sugar metabolism across all lines. These findings are also mirrored when we use Ribo-seq data for the KEGG analysis (Fig. S9A).

**Figure 6:**
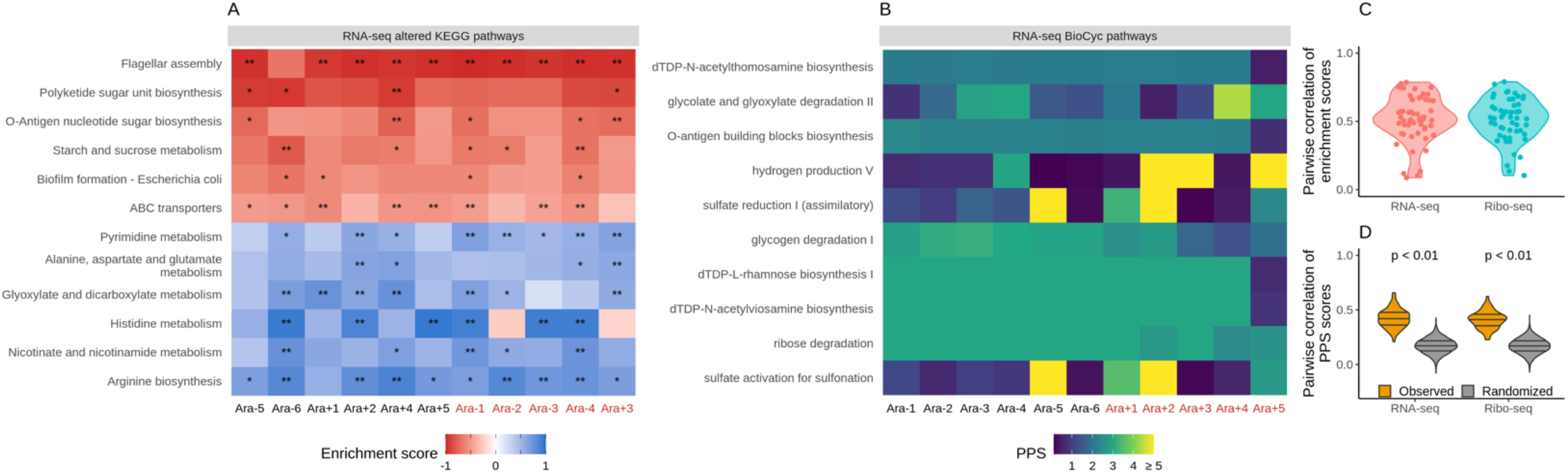
**A**. Parallel changes in functional categories. KEGG enrichment scores from the RNAseq data. Enrichment score represents the degree to which a pathway was up (positive) or downregulated (negative). The functional categories are ordered by increasing the mean enrichment score across the lines. **B**. Pathway perturbation score (PPS) is calculated from RNA-seq fold changes. Higher PPS indicates larger degrees of alteration but does not indicate directionality. **C**. Pairwise correlations of KEGG enrichment scores for all pathways that were significantly altered in at least one line. **D**. Pairwise correlations of PPS scores. PPS scores for the randomized set was calculated by randomizing the fold-changes within each line.

**Figure 7.**
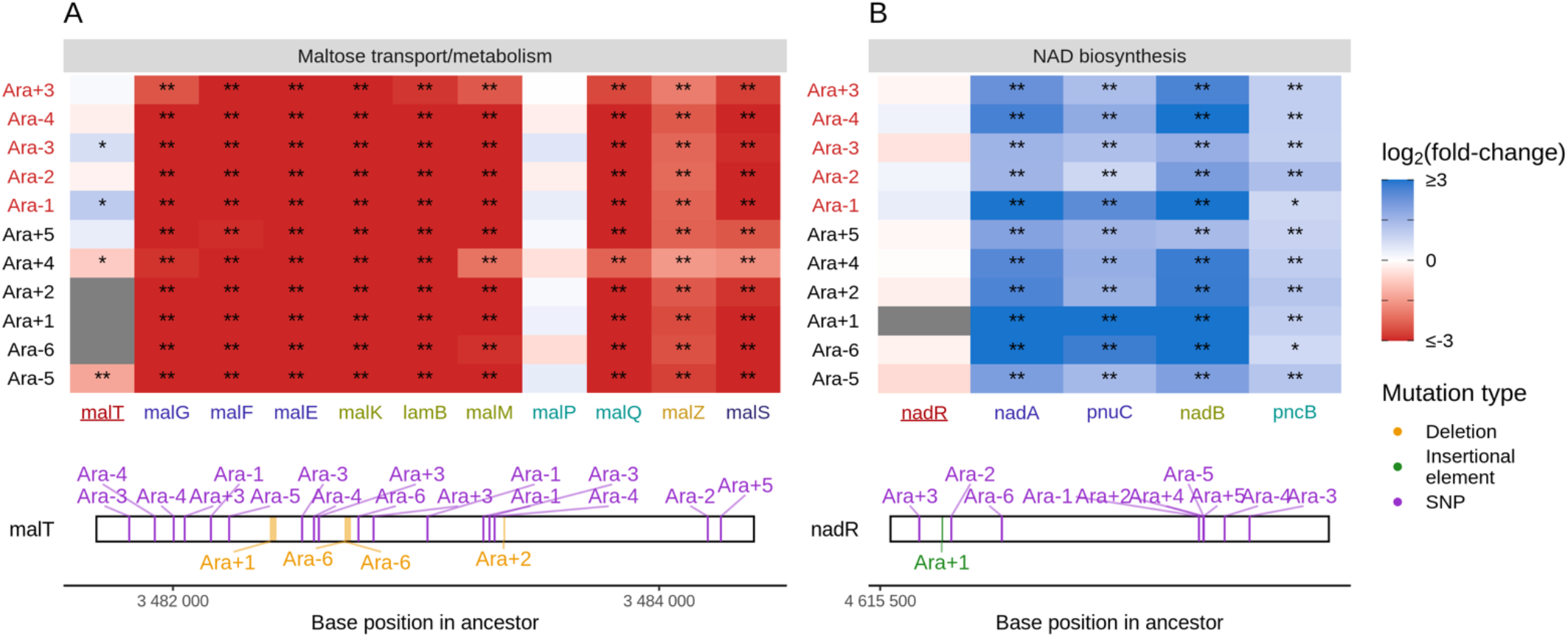
Mutations in transcriptional regulators lead to parallel changes in gene expression. RNA-seq fold-changes for genes belonging to **A.** maltose-transport/metabolism and **B.** NAD biosynthesis. Gene names in each category are colored based on their operon membership. Mutations in transcriptional activator malT decrease expression of its downstream genes/operons. Mutations in transcriptional repressor nadR increase expression of its downstream genes/operons. Asterisks indicate statistical significance of fold-changes, ** q ≤ 0.01, * q ≤ 0.05. Grey panels in the heatmap indicate gene deletion. Lower panels show the type and location of mutations in each transcription factor.

While KEGG pathway analysis encompasses molecular interactions and reaction networks, we wondered which specific metabolic pathways were altered across all lines and which ones remained mostly unchanged over 50,000 generations. Because *E. coli* REL606 is annotated in the Biocyc collection of databases, we used their metabolic mapping tool to score pathway alterations with a pathway perturbation score (PPS) in each of the evolved lines (see methods for a detailed explanation of the scoring). Similar to the KEGG pathway analysis, we find a high degree of parallelism, even at the level of specific metabolic pathways (Fig. 6B, D). Interestingly, 4 out of 5 most altered pathways are involved in lipopolysaccharides (LPS) biosynthesis, a major component of Gram-negative bacteria’s outer membrane. This indicates that in addition to changes in cell size and shape, the composition of the evolved lines’ outer membrane has significantly changed. Nonetheless, there is a core set of unaltered pathways, even in clones with a mutator phenotype. Pathways with low PPS scores, indicating low levels of alteration included D-serine degradation (mean RNAseq PPS = 0.12, sd = 0.13), pseudouridine degradation (mean RNAseq PPS = 0.11, sd = 0.06), and others (see Table S12 for complete PPS scores). These may represent pathways with activity levels that cannot be altered or whose alteration provides little to no fitness benefit.

### Mutations to transcriptional regulators explain many parallel expression changes

Given the high degree of parallelism in evolved lines at the gene expression level, we wondered whether some of these patterns could be explained by a parallel set of mutations at the genetic level. Because KEGG, PPS, and GO analyses all identified metabolism and catabolism of various sugars to be significantly altered, we started by looking at mutations to genes involved in these categories. Previous work has shown that depending on the generation sampled, evolved clones grow poorly (20,000th generation) or not at all (50,000th generation) on maltose^45^. Because maltose is absent from the growth media in the LTEE, maintenance of these transporters is likely unnecessary^46^. Additionally, at 20,000 generations, the transcriptional activator of the operon responsible for maltose metabolism, malT, was the frequent target of mutations that reduced its ability to act as a transcriptional factor, and introduction of malT mutations in the ancestor had a fitness benefit^46^. In *E. coli*, MalT regulates the transcription of several operons - *malEFG* (maltose ABC transporter), *malK*-*lamB*-*malM* (MalK, part of maltose ABC transporter; LamB, maltose transporter; MalM, conserved gene of unknown function, MalPQ (two enzymes involved in maltose metabolism), and the genes *malZ* (maltodextrin glucosidase) and *malS* (an α-amylase). We find that each of these operons was consistently and significantly downregulated across all lines (Fig. 6E). Changes to the LamB transporter have also been shown to affect susceptibility to phage infection in the LTEE^47^.

In the LTEE, NadR, a transcriptional repressor of genes involved in NAD biosynthesis, is known to be frequently mutated, with many mutations occurring in its DNA binding domain ^48,49^. In fact, all evolved clones used in this study are known to have some mutation in *nadR*^12^. Given the high frequency of parallel inactivating mutations in *nadR*, it is likely that these mutations are adaptive as they might increase intracellular NAD concentrations leading to faster growth^48,49^. We find that genes directly under the regulation of *nadR* -- the *nadAP* operon consisting of *nadA* (quinolinate synthase) and *pnuC* (nicotinamide riboside transporter), and genes -- *nadB* (L-aspartate oxidase) and *pncB* (nicotinate phosphoribosyltransferase, were significantly upregulated in all lines. We also found enrichment of NAD pathways based on KEGG, GO (GO:0019674, GO:0009435), and PPS analysis. Interestingly, four non-operonic genes *nadCDEK*, which play various NAD biosynthesis roles but are not regulated by *nadR*, were largely unaltered (*nadE* was statistically significantly upregulated in 4 lines, DESeq2 q ≤ 0.01, Table S7). Concordantly, their transcriptional regulator, *nac*, is rarely mutated. This may suggest some sort of specificity to how NAD levels may be increased.

In addition to linking the effects of specific mutations on gene expression changes in maltose and NAD regulation, we have also identified mutations that likely change the expression of genes involved in arginine biosynthesis, glyoxylate bypass system, and copper balance (Fig. S11, see supplementary methods). However, there also exist several functionally-related sets of genes, such as flagellar assembly, sulfur homeostasis, and biosynthesis of one-carbon compounds – that have parallel changes in expression levels without any obvious sets of parallel mutations linking these changes (Fig. S11). The data generated in this study will likely prove to be a rich resource for understanding the metabolic changes that occur over long periods of evolution in a simple environment such as in the LTEE, thereby adding a rich new dimension to the well-studied mutational changes and gene-expression changes described here.

## Discussion

Adaptation to novel environments often takes unique mutational paths even when the tempo and mode of adaptation are similar across populations^8,12,50–53^. This is due, in part, to the fact that most genetic networks are highly redundant and that many mutations have pleiotropic effects. To bridge the gap between parallel fitness gains in a system with mostly unique genetic changes, we wanted to study gene expression – a main link between genotype and fitness. To that end, we generated RNA-seq and Ribo-seq datasets for individual clones from the ancestral strains and 11 populations evolving under a constant environment for 50,000 generations in the *E. coli* long-term evolution experiment. Using these datasets, we have characterized the landscape of gene expression changes and elucidated several key features of the molecular mechanisms involved. First, we show that the evolved lines in the LTEE have remarkably parallel exponential phase expression profiles after 50,000 generations. Second, these changes primarily occurred at the transcriptional level, with translational changes following suit. Nonetheless, we identified signatures of global increases in translation termination rates. Third, transcriptional regulators of genes that were mutated in multiple lines had similar functional effects on their downstream targets across all lines. This indicates a strong penetrance of mutational effects to the phenotypic level even when half of the evolved lines had a hypermutable phenotype. Fourth, we show how functional consequences of mutations are consistent with adaptation in a constant environment -- genes involved in central metabolism and amino-acid biosynthesis are consistently upregulated, and genes involved in sensing environmental changes and stress responses are downregulated.

Relating gene expression changes to specific mutations in LTEE is far from perfect. For many genes that are functionally related and show parallel changes in gene expression, such as the ones involved in flagellar assembly and sulfur homeostasis, we find few mutations around their coding sequences or sequences of their known transcriptional regulators. This might be due to two factors: (i) a lack of complete knowledge of gene regulatory networks underlying these functions, and (ii) parallel epigenetic changes such as changes in DNA supercoiling heterogeneities, affecting promoter activity^54^. Indeed, changes to DNA superhelicity occur in multiple LTEE lines^55^. Another key challenge in attributing expression changes to mutations is that half of the evolved lines in LTEE have a hypermutable phenotype. These genotypes have ~100-fold higher mutational load than their non-mutator counterparts. It is remarkable that despite a higher mutational burden, expression patterns between mutator and non-mutator lines are highly correlated, suggesting that the bulk of the additional mutations are indeed passenger mutations^13^. While our current study has focused on expression patterns in the exponential phase, populations in the LTEE spend most of their time before serial transfer in the stationary phase. However, it remains unclear if we would observe a similar level of parallelism in the stationary growth phase or how similar the expression profiles might be across distinct growth phases. Taking a multi-omics approach, like the one presented above, will provide critical insights into the tradeoff between expression patterns across phases. Lab evolution experiments combined with high-throughput multi-level sequencing approaches offer a rich resource for studying the molecular mechanisms underlying complex adaptations and provide insights into the repeatability of evolution.

## Supporting information

Supplemental Table 1 - read counts

Supplemental Table 2 - 3nt periodicity

Supplemental Table 3 - cell size measurements

Supplemental Table 4 - colony count data

Supplemental Table 5 - ERCC molecules per sample

Supplemental Table 6 - molecules per CFU

Supplemental Table 7 - expression fold-changes

Supplemental Table 8 - mutations

Supplemental Table 9 - riborex results

Supplemental Table 10 - ribosome-densities

Supplemental Table 11 - KEGG results

Supplemental Table 12 - PPS scores

Supplemental Table 13 - GO enrichment results

## ACKNOWLEDGEMENTS

We thank Richard Lenski for generously providing clones from ancestral and 50,000 generations of the LTEE. P.S. is supported by NIH/NIGMS grant R35 GM124976, NSF DBI 1936046, subcontracts from NIH/NIDDK R01 DK056645, R01 DK109714, and R01 DK124369, as well as start-up funds from the Human Genetics Institute of New Jersey at Rutgers University. S.S.Y is supported by start-up funds from the Waksman Institute and Rutgers University

## AUTHOR CONTRIBUTIONS

P.S. conceived the study and designed the experiments; J.S.F., S.L, and S.S.Y. conducted experiments; J.S.F., S.S.Y., and P.S. analyzed data. J.S.F. and P.S. wrote the manuscript with input from S.L. and S.S.Y.

## METHODS

### Bacterial cell culture, recovery, and lysis

Richard Lenski generously provided clones from LTEE. Specifically, the following clones were used: Ara-1, 11330; Ara+1, 11392; Ara-2, 11333; Ara+2, 11342; Ara-3, 11364; Ara+3, 11345; Ara-4, 11336; Ara+4, 11348; Ara-5, 11339; Ara+5, 11367; Ara-6, 11389; Ara+6, 11370. Clones were grown in DM25 medium (HiMedia M390) supplemented with 4 g/L glucose. Each culture was grown in 50 mL in a shaking incubator at 37 C at 125 rpm until an OD600 of 0.4-0.5 was reached. Cells were recovered via vacuum filtration and immediately frozen in liquid nitrogen (LN_2_). Frozen pellets were stored at −80 C until lysis. For lysis, a mortar and pestle were chilled to cryogenic temperatures with LN_2_. The pellet was ground to a powder while submerged in LN_2_. Once pulverized, 650 uL of lysis buffer was added to each sample and ground further. Lysis buffer contained the following: 20 mM Tris pH 8, 10 mM MgCl_2_, 100 mM NH_4_Cl, 5 mM CaCl_2_, 1 mM chloramphenicol, 0.1% v/v sodium deoxycholate, 0.4% v/v Triton X-100, 100 U/mL DNase I, 1 uL/mL SUPERase-In (Thermo Fisher Scientific AM2694). The frozen lysate was allowed to thaw until liquid, then incubated for 10 min on ice to allow complete lysis. Afterward, the lysate was centrifuged at 20,000g for 10 minutes at 4 C, and the supernatant recovered and transferred to a new tube. Each sample was split into two for RNA-seq and Ribo-seq libraries.

### RNA-seq library preparation

Lysate destined for RNA-seq libraries was subjected to total RNA extraction using the Trizol method (Thermo Fisher Scientific 15596026) as per the manufacturer’s instructions. RNA was quantified using UV spectrophotometry. We used the ERCC RNA Spike-In Mix (Thermo Fisher Scientific 4456740) in library preparation. For RNA-seq libraries, 3 uL of a 1:100 dilution of the set 1 oligos was added to the first replicate and 4 uL to the second replicate. The spike-ins were added directly to the lysate destined for RNA-seq before Trizol based RNA extraction. 2 ug of RNA with ERCC controls were subjected to fragmentation in a buffer containing final concentrations of 1 mM EDTA, 6 mM Na_2_CO_3_, and 44 mM NaHCO_3_ in a 10 uL reaction volume for 15 minutes at 95 C. 5 uL of loading buffer (final concentrations of 32% v/v formamide, 3.3 mM EDTA, 100 ug/mL bromophenol blue) was added to each sample, and the resulting 15 uL mixture was separated by gel electrophoresis with a 15% polyacrylamide TBE-urea gel (Invitrogen EC68852BOX) at 200 V for 30 minutes. Gels were stained for 3 minutes with SYBR Gold (Thermo Fisher Scientific S11494), and the region corresponding to the 18-50 nucleotide sized fragments excised. We excised this region so that we would have similarly sized fragments for both RNA-seq and Ribo-seq libraries. RNA was recovered from the extracted fragments by adding 400 uL a buffer containing 300 mM sodium acetate, 1 mM EDTA, and .25% w/v SDS, and freezing the samples on dry ice for 30 minutes. Then, samples were incubated overnight on a shaker at 22 C. 1.5 uL of GlycoBlue (Thermo Fisher Scientific AM9515) was added as a co-precipitant, followed by 500 uL of 100% isopropanol. The samples were chilled on ice for 1 hour then centrifuged for 30 minutes at 20,000g at 4 C. The supernatant was removed, and the pellet was allowed to air dry for 10 minutes. The pellet was resuspended in 5 uL of water, and 1 uL was used to check RNA concentration via UV spectrophotometry.

### Ribo-seq library preparation

Lysate destined for Ribo-seq was incubated with 1500 units of micrococcal nuclease (Roche 10107921001) and 6 uL of SUPERase-In at 25 C for 1 hour and shaken at 1400 rpm. 2 uL of .5 M EGTA pH 8 was added to quench the reaction, which was then placed on ice. The reaction was centrifuged over a 900uL sucrose cushion (final concentrations of 20 mM Tris pH 8, 10 mM MgCl_2_, 100 mM NH_4_Cl, 1 mM chloramphenicol, 2 mM DTT, .9 M sucrose, 20 U/mL SUPERase-In) using a Beckman Coulter TLA100 rotor at 70,000 rpm at 4 C for 2 hours in a 13 mm x 51 mm polycarbonate ultracentrifuge tube (Beckman Coulter 349622). The sucrose solution was removed from the tube, and the pellet resuspended in 300 uL of Trizol, mixed by vortexing, and RNA was extracted according to the manufacturer’s protocol. Samples were then separated by gel electrophoresis and purified in the same manner as for RNA-seq.

### Unified library preparation

Once fragments were obtained from RNA-seq and Ribo-seq samples, they could be subject to a unified library preparation protocol. In total, 8 pooled libraries were prepared, with each library consisting of a single replicate of 6 Ara+ or 6 Ara-clones of one sequencing type. For example, one library would consist of replicate 1 of Ara-1-6 for RNA-seq, and another would consist of the second replicate. The final library structure was 5’ adapter - 4 random bases - insert - 5 random bases - sample barcode - 3’ adapter. The randomized bases function as UMIs for deduplication.

3’ dephosphorylation was performed by incubating fragments with 10 U/uL T4 Polynucleotide Kinase (New England Biolabs M0201S) in the supplied buffer (NEB B0201S) along with SUPERase-In for 1 hour at 37 C in a reaction volume of 5 uL.

Linker ligation took place by adding the following reagents to the above reaction to the indicated final concentrations: 17% w/v PEG-8000, 200 U/uL of T4 RNA Ligase 2 (NEB M0351S), 1X T4 RNA Ligase Reaction Buffer (NEB B0216L), and 20 uM pre-adenylated linkers. The reaction volume totaled 10 uL, and was incubated for 3 hours at 22 C. Afterwards, 10 U/uL of 5’ deadenylase (NEB M0331S), 10 U/uL Rec J exonuclease (Epicentre RJ411250), and the included buffer were added and incubated at 30 C for 45 minutes.

RNA was purified using a Zymo Research Oligo Clean & Concentrator Kit (Zymo, D4060), and then rRNA depleted using the Illumina Ribo-Zero rRNA Depletion Kit for bacteria, both steps being performed according to the manufacturer’s instructions.

5’ phosphorylation was performed by mixing 6 uL of rRNA depleted RNA with 1 uL of 10X PNK buffer (NEB B0201S), 1 uL of PNK enzyme (NEB M0236S), and 2 uL of 1mM ATP to total 10 uL and incubated at 37 C for 30 minutes followed by inactivation by heating to 65 C for 20 minutes.

Hybridization with the reverse transcription primers was performed by adding 1 uL of SR RT Primer (NEB E7333A) to the above reaction and incubating at 75 C for 5 minutes, 37 C for 15 minutes, and 25 C for 15 minutes.

5’ adapter ligation was performed by adding 3 uL of 10uM 5’ adaptor (which was previously denatured by heating to 70 C for 2 minutes and placed on ice, NEB E7330L), 2 uL of 10X T4 RNA ligation buffer (NEB B0216L), 2 uL of 10mM ATP, 2 uL of T4 RNA ligase I (NEB M0204S) totaling 20 uL and incubated for 1 hour at 30 C.

Reverse transcription was performed by adding the following to the above reaction: 8 uL of 5x first strand buffer (NEB E7330L), 2 uL of 10mM dNTPs (each), 4 uL of 10X DTT (Invitrogen *something*), 2 uL of SUPERase-In, 2uL of SuperScript II (NEB M0368L), and 2 uL of water, totaling 40 uL and incubated at 50 C for 1 hour then inactivated by heating to 70 C for 15 minutes.

PCR amplification of the above reaction was performed by taking 150 ng of cDNA template and adding 10 uL 5X Phusion HF buffer (Thermo Fisher Scientific F518L), 1uL 10 mM dNTPs (each), 1.25 uL 10uM SR primer (from NEB E7330L), 1.25 uL 10uM index 3 primers, .5 uL of Phusion polymerase (NEB M0530S), and enough to water to total the reaction volume at 50 uL. This was cycled as follows in a thermocycler: 30 sec at 90 C; 14 cycles of 15 sec at 94 C, 30 sec at 62 C, 15 sec at 70 C; 5 min at 70 C.

PCR products were separated by gel electrophoresis on a 6% polyacrylamide gel at 120 V for 45 minutes. The region corresponding to the expected product size was excised and purified from the gel by soaking the resected pieces in 250 ul DNA gel elution buffer (NEB E7324A) at 22 C and 200 rpm overnight on a rotator and transferring the solution to a gel filtration spin column (Corning 8160) and centrifuging for 2 minutes at 16,000g. 1.5 uL of GlycoBlue, 25 uL of 3M sodium acetate pH 5.5, and 750 uL of 100% ethanol were added, and the solution was held on ice for 2 hours, then centrifuged at 20,000g at 4 C for 30 minutes. The supernatant was removed, and the pellet washed with 75 % ethanol and again centrifuged at 20,000g at 4 C for 5 minutes. The pellet was allowed to air dry and resuspended with 11 uL of water. 1 uL was used to check concentration via UV spectrophotometry. The completed libraries were sequenced on Illumina NextSeq in 75 bp single-end mode.

### ERCC spike-in controls and modeling

ERCC RNA Spike-In Mix (Thermo Fisher Scientific 4456740) was used in library preparation. For RNA-seq libraries, 3 uL of a 1:100 dilution of the set 1 oligos was added to the first replicate and 4 uL to the second replicate. The spike-ins were added directly to the lysate destined for RNA-seq before Trizol based RNA extraction. The file “absolute_counts.Rmd” contains the code for the linear modeling using the ERCC data.

### CFU determination

Before recovery, 1mL of culture was extracted for CFU determination. LB agar plates were used for colony growth. We performed a dilution series of that 1mL culture from 1:10 to 1:1e6 in increments of 10. 100uL of each dilution was spread on a plate and incubated overnight at 37C. We determined CFU counts manually from the most appropriate dilution for each culture, usually between 1:1e3 and 1:1e6 dilutions.

### Optical microscopy

#### (i) Media and growth conditions

Liquid cultures were grown at 37°C with aeration, unless otherwise indicated, in DM25 liquid medium (Davis minimal broth supplemented with glucose at a concentration of 25 mg per L^10^).

#### (ii) Microscopy

Prior to each experiment, clones were grown in liquid cultures in DM25 medium overnight at 37°C with aeration. OD_600_ of the cultures were ∼0.1–0.3. Microscope slides were prepared with 1% agarose pads, and cells were imaged by microscopy. Phase-contrast microscopy was performed using an Olympus IX81 microscope with a 100-W mercury lamp and 100× NA 1.35 objective lens. 16-bit images were acquired with a SensiCam QE cooled charge-coupled device camera (Cooke Corp.) and IPLab version 3.7 software (Scanalytics) with 2 × 2 binning. Average cell lengths were determined from phase contrast images using ImageJ^56^ and the MicrobeJ plugin^57^.

### Sequencing data processing

Sequencing data are deposited here - https://www.ncbi.nlm.nih.gov/geo/query/acc.cgi?acc=GSE164308. Code for all data processing and subsequent analysis can be found in a series of R markdown documents here – (https://github.com/shahlab/LTEE-gene-expression). The file titled “data_processing.Rmd” contains the code for processing of the raw sequencing data. We processed 8 raw data files. We used Cutadapt^58^ to remove adapters and retained only reads that had successful trimming. We then used the dedupe.sh script from the BBtools suite to remove PCR duplicates. Files were demultiplexed using the FASTX-Toolkit barcode splitter script. After demultiplexing, barcodes and the randomized adapters were removed using cutadapt. The 4 nucleotide UMIs were removed from the 5’ end of a read and 10 nucleotides from the 3’ end (5 UMI + 5 barcode). Only reads longer than 24 nucleotides after trimming were retained.

### Alignment

#### Differential expression

Code for this section can be found in the file titled “DEseq2.Rmd”. We used DEseq2^30^ with the “apeglm” normalization^59^ for differential expression. In estimating fold-changes, we compared the 4 replicates of the ancestors (2 each from ancestors of Ara+ and Ara-) to 2 replicates of each of the evolved lines. Because some genes in some lines contained indels or were deleted entirely, some transcripts were missing from the transcriptome fastas used to create indices for alignment. We added these genes back to Kallisto’s counts with estimated counts of 0 and assigned them fold-changes of NA. Count matrices containing identical complements of transcripts were used in the differential expression analysis for each line, such that all evolved lines had the same complement of genes as the ancestors.

#### Change in ribosomal density analysis

We used Riborex^34^ to analyze changes in ribosomal density. The same count matrices used for DEseq2 were used here, and comparisons were made in the same manner of 4 ancestral samples (2 lines, 2 replicate each) to 2 evolved clones (1 line, 2 replicates). The code for this section can be found in the file “riborex.Rmd”

#### Codon specific positioning of Ribo-seq data

Code for this section can be found in the file “codon_specific_densities.Rmd”. We used hisat2^60^ to align our Ribo-seq data to each clone’s unique genome and marked the A site position of a read using a fixed offset of 37nt from the 3’ end of a read. It has been shown that mapping bacterial Ribo-seq reads by their 3’ ends is more accurate than 5’ mapping^61^. We then calculated genome-wide ribosome density at each codon using only genes that had at least 100 reads. The distributions of read counts per gene can be seen in figure S1C. Only bacterial protein-coding genes (not tRNA or insertional element genes) were considered. To calculate ribosome densities on a codon for a gene, the number of reads mapping to a codon was normalized to the total number of reads mapping to that gene in a replicate and line-specific manner. Genome-wide codon density is calculated by taking genes with at least 100 reads mapping to them and taking the average number of normalized reads mapping to each codon across that set of genes as the genome-wide codon density.

#### Functional analysis

We used three different functional analysis methods – GO (using the R package topGO), KEGG (using the R package clusterprofiler^62^, and PPS^44^. The code for each of these analyses can be found in the Rmd files named “go. Rmd”, “kegg_analysis.Rmd”, and “manual_PPS.Rmd,” respectively. We used a manual implementation of the Biocyc PPS score because the website was not capable of high throughput analysis. Briefly, each pathway is composed of at least one reaction, and each reaction is completed by at least one enzyme. First, a reaction perturbation score is calculated for each reaction in a pathway. It is defined as the absolute value of the largest fold-change of an enzyme associated with that reaction. To calculate PPS, for a pathway having N reactions, PPS = sqrt((Σ RPS^2^) / N).

### SUPPLEMENTAL ANALYSIS

#### Cell size and filamentation

Evolved lines form filaments more frequently and form longer filaments compared to the ancestor. This is supported by the fact that all evolved lines except Ara+1 had significantly longer cells compared to the ancestor (Welch’s t-test, p<.0001 for all lines) (Fig. S2A). Additionally, volume and aspect ratio are positively correlated in all lines (0.53 ≤ R ≤ 0.94). Length and volume was positively correlated (0.76 ≤ R ≤ 0.95), but width and volume showed a low correlation (0.12 ≤ R ≤ 0.45) (Fig. S2C). Taken together, increases in the volume to large values are due to increases in one dimension, length, suggesting increased filamentation. We designated cells that are greater than three times the median volume of a given line as filaments. Even after removing filaments from the comparisons, each evolved line was still larger in volume than the ancestor (Fig. S2B). Removal of filaments did not alter the relationship between the median volume and RNAs per CFU (Fig. S2D).

#### CFU counts

One caveat to the relationship between CFU counts and RNA abundance is that the CFU counts may be misleading, especially in light of the increased filamentation suggested by our microscopy data. Because a single chain of bacteria composed of multiple cells could be the source of a single colony, the CFUs may be an underestimate of the number of cells that had gone into the preparation of each of the evolved lines libraries. If this was the case, it might contribute to the observed results.

#### GO analysis

We also performed GO searches in all three ontologies, Cellular compartment (CC), Biological process (BP), and Molecular function (MF). The top 5 up and downregulated terms for each ontology can be seen in Fig. S11, and the complete results can be found in Supplementary Table S13. These searches found results similar to the KEGG and PPS results. For example, terms related to the flagellar apparatus (BP, GO:0044780, GO:0044781, GO:0071978, GO:0097588, GO:0071973, GO:0001539; CC, GO:0009288 GO:0009424, GO:0044461), polysaccharide transport (BP, GO:0015774, GO:0033037), specifically, maltodextrin transport (BP, GO:0042956), arginine biosynthesis (BP: GO:0006526), and others reach statistical significance (Fisher’s exact test, p ≤ 0.05) in many of the lines. Other terms related to iron were also found to be enriched and many genes related to iron transport or incorporation into organic molecules were found to have significant fold-changes in the DESeq2 results (data not shown, see table S7 for complete DESeq2 results).

#### Analysis of altered pathways

Flagella are used for bacterial motility and allow bacteria to move to new environments by swimming. Previous experiments in the LTEE have shown the downregulation of flagellar apparatus genes in Ara+1 and Ara-1 at 20,000 generations, though the exact source of these downregulations was not determined^16^. We find that genes related to the flagellar apparatus are significantly downregulated in 10 of the 11 lines considered here (Fig. S11A). The flgBCDEFGHIJK, flgAMN, and flhABE operons are significantly downregulated in all but Ara-6, where only some of these genes were downregulated. These operons contribute various proteins to the flagellar apparatus and are regulated in part by the transcription factors flhC and flhD, which have complicated regulation dictated by various environmental factors^63^. flhC and flhD are downregulated in 3 of the evolved lines but mostly unaltered in the others. These genes are rarely mutated in the clones used in this study (Fig. S11A, bottom). The fitness benefits of downregulation to the flagellar apparatus may be multifaceted. The flagellar apparatus is an expensive piece of machinery to produce, and it requires energy to move. Other *E. coli* evolution studies have shown that mutations in flagellar genes are common and provide a fitness advantage^64^. Additionally, the *E. coli* B strain is thought to be non-motile^65^. Taken together, the downregulation of flagella may simply be the removal of an unused system. Surprisingly, the lack of parallel changes in transcriptional regulators flhCD indicates that it is unlikely that transcriptional changes are the primary mode for downregulation of the flagellar protein operons.

Amino acids are the building blocks for proteins, and translation of new proteins is required for cellular growth. Hence, increased levels of intracellular amino acids would allow faster translation of proteins and faster growth. Terms involving amino acid biosynthesis showed up frequently in all three methods used for functional analysis (KEGG, GO, and PPS). Arginine biosynthesis (KEGG and GO:0006526) was a frequently upregulated category. We find that genes related to arginine biosynthesis were upregulated in 8 out of 11 lines (Fig. S11B). These genes are partly controlled by the argR repressor, which represses their transcription when L-arginine is abundant^66^. 5 out of 10 lines had mutations to the argR coding sequence, and other lines had mutations occurring nearby. Interestingly, we find that expression levels of argR remain unchanged in all lines indicating that these mutations may have disabled argR function, causing de-repression of its downstream targets.

The glyoxylate bypass system allows *E. coli* to utilize acetate as a carbon source, is composed of the aceBAK operon, and regulated by iclR and arcAB^67^. Acetate is a metabolic by-product but can be returned to central carbon metabolism for biosynthetic reactions by this system. Previous studies have shown that mutations in iclR and arcB cause depression of their target genes are beneficial in the LTEE^68^. Consistent with these results, we found that the aceBAK operon was upregulated in 9 of 11 evolved lines (Fig. S11C).

Copper and silver have antibacterial properties^69^, and bacteria have evolved systems to mitigate toxicity from these elements. The cusCFBA operon, regulated by the cusRS sensor kinase, codes for proteins that transport copper and silver ions out of the cell^70^. Additionally, the cytoplasmic copper chaperone copA, regulated by cueR^71^, and cueO (multicopper oxidase^72^) regulate copper homeostasis in the cell. These genes contained deletions 5 of our clones and were downregulated in 3 of the 6 lines where they remained (Fig. S11D). Overall, 8 of the 11 lines surveyed here had defects in these systems. This suggests that there may be the selection for removal or downregulation of these genes. In contrast to natural environments, the laboratory environment is likely free of copper and silver, rendering these systems dispensable.

Sulfur is a critical component of many biological molecules, like amino acids, and participates in creating other structures like iron-sulfur cluster proteins. Organic sulfur is transported across the cell membrane by proteins from the cysPUWAM operon, which encodes for a sulfate/thiosulfate importer^73^, the gsiABCD operon which encodes for a glutathione importer^74^, the tauABCD operon which codes for a taurine importer^75^, and tcyP, the major L-cysteine importer^76^. We found that many of these genes were downregulated in many of the lines (Fig. S11E). The cysB gene positively regulates these genes and was downregulated in most lines. This gene contained few mutations across the lines. The sources of organic sulfur in the medium used in the LTEE are ammonium and magnesium sulfate, for which the cysPUWAM operon functions as the importer. The mechanism and reasons for alterations to these operons remain unclear. The amount of organic sulfur in the medium may be sufficient to allow the downregulation of sulfur transport systems without impacting downstream pathways that require sulfur.

Glycine plays a role in protein construction and can serve as a building block for other metabolic pathways such as one-carbon metabolism or serine synthesis^67,77^. We found that the gcvTHP operon, which encodes for proteins in the glycine cleavage system, were upregulated in 6 of the 11 lines. Increases in the levels of compounds involved in this set of reactions directly may increase the growth rate. Though there are some mutations in and around transcriptional regulators of these genes, their effects are unclear. Whether changes to these genes are due to changes in their transcription factors or other changes, the upregulation of these genes in many lines suggests that it may be beneficial.

## Supplemental tables

A description of the supplemental tables:

Table S1: The file “table_s1_read_counts.csv” contains quantification of read counts per gene based on Kallisto for each sample. Counts in this file were rounded, and new TPMs were calculated based on rounded counts. This file was generated using “data_cleaning.Rmd”.

Table S2: The file “table_s2_three_nt_periodicity.csv” contains the data needed to show periodicity in the Ribo-seq data. This file was generated using “3nt_periodicty.Rmd”.

Table S3: The file “table_s3_cell_size.csv” contains cell size data derived from phase-contrast microscopy.

Table S4: The file “table_s4_colony_counts.csv” contains information about colony forming units for each of the samples.

Table S5: The file “table_s5_ercc_molecules_per_sample.csv” contains information about the use of ERCC controls in each sample and the counts/TPMs of each control in each sample.

Table S6: The file “table_s6_mRNAs_per_cfu.csv” shows the absolute counts of mRNAs per CFU for each gene in each sample. This file was generated using “absolute_counts.Rmd”.

Table S7: The file “table_s7_fold-changes.csv” contains the results of gene-expression fold-changes based on DESeq2 analysis. This file was generated using “DEseq2.Rmd”.

Table S8: The file “table_s8_mutations.csv” contains data on mutations accumulating in LTEE and was derived from Good et al. 2017 and downloaded from https://barricklab.org/shiny/LTEE-Ecoli/.

Table S9: The file “table_s9_riborex_results.csv” contains the results of differential ribosome-density analysis using Riborex. This file was generated using “riborex.Rmd”.

Table S10: The file “table_s10_genome_wide_codon_densities.csv” contains the genome-wide codon-specific ribosome-densities. This file was generated using “codon_specific_densities.Rmd”.

Table S11: The file “table_s11_kegg_results.csv” shows the results of KEGG enrichment analyses. This file was generated using “kegg_analysis.Rmd”

Table S12: The file “table_s12_pps_scores” shows the PPS scores analyses. This file was generated using “manual_PPS.Rmd.”

Table S13: The file “table_s13_go_results.csv” shows the results of GO enrichment analyses. This file was generated using “go.Rmd”.

---

**Figure S1:**
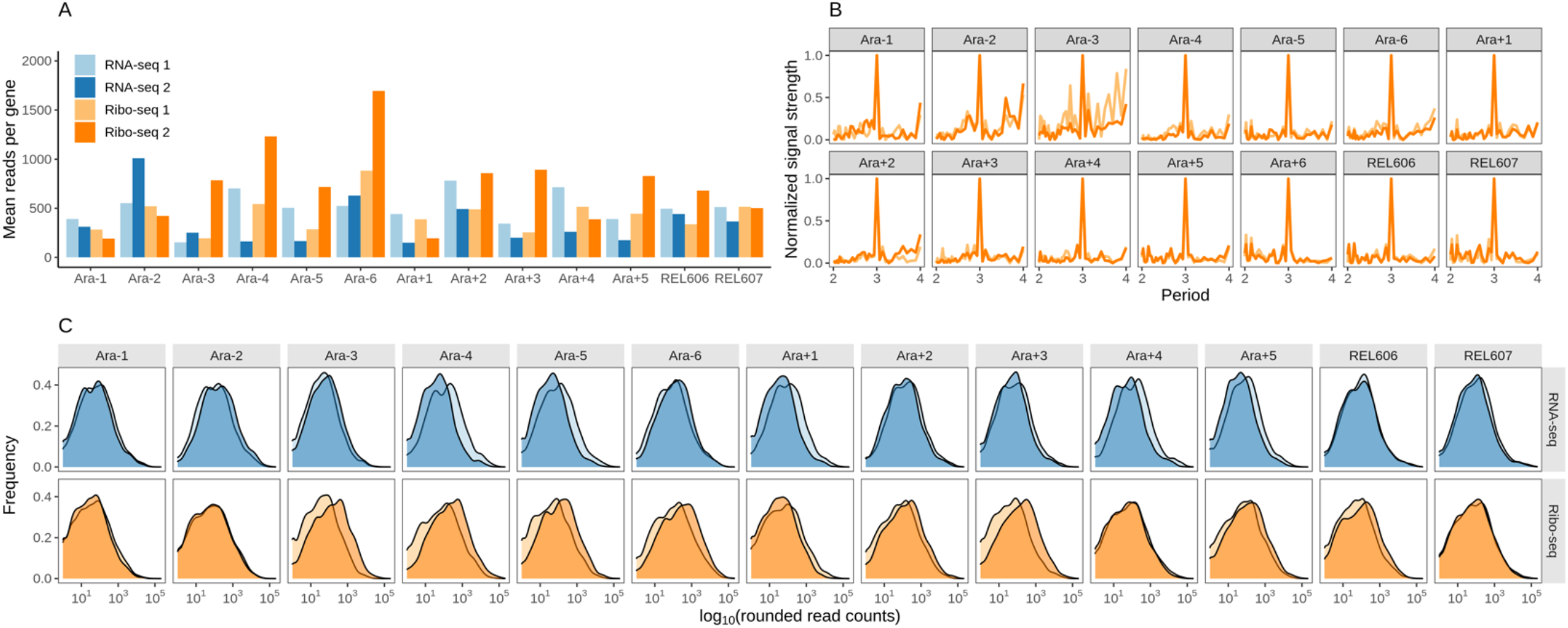
Summary of sequencing data. **A.** The average number of reads aligned per protein-coding gene by Kallisto for each sample. The color scheme remains the same for the other panels. **B.** The periodicity of the ribo-seq datasets is determined using a fast Fourier transform (see methods). **C.** Distributions of reads per protein-coding gene in each sample.

**Figure S2:**
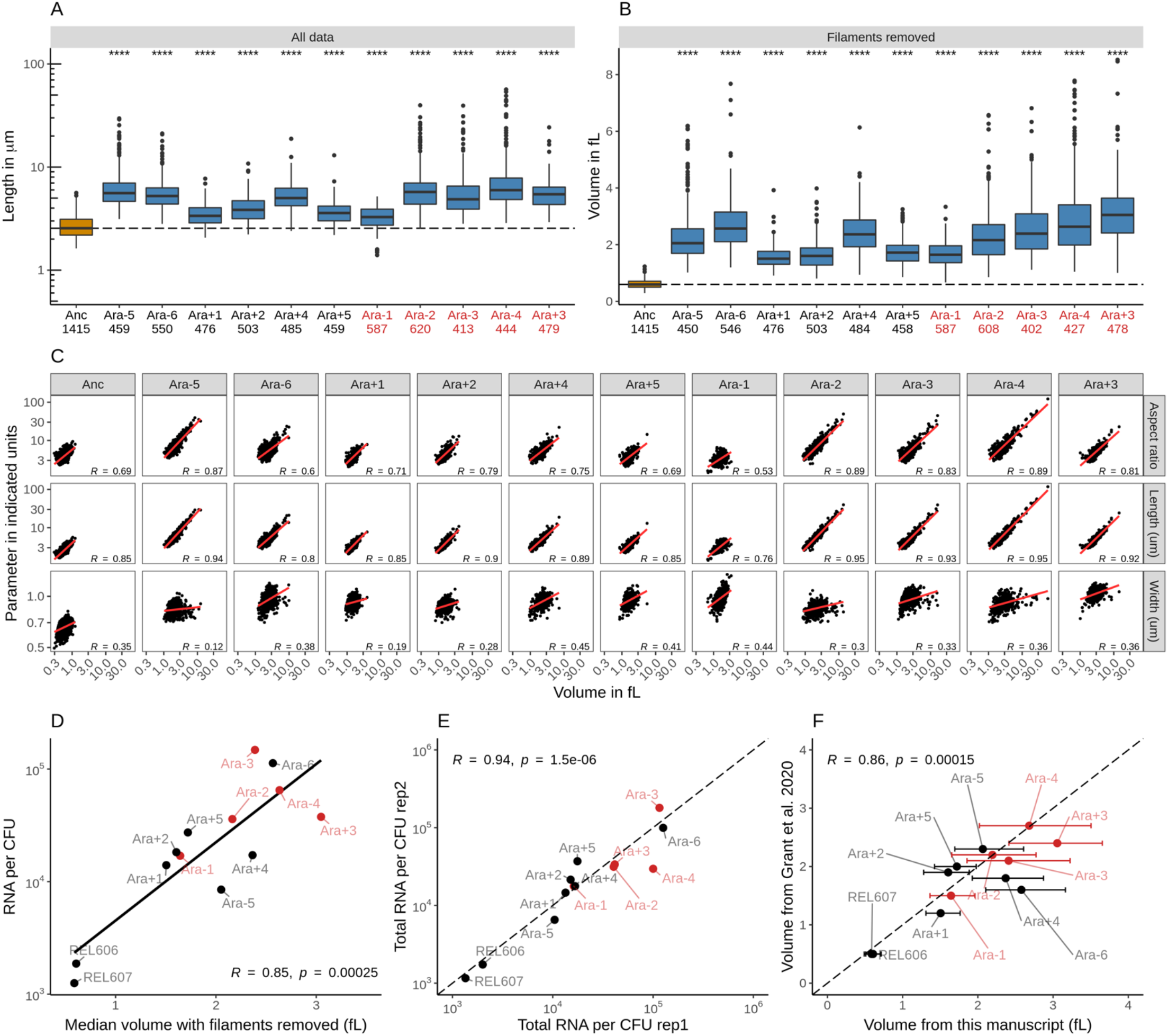
**A**. Length distributions of cells as determined by phase contrast microscopy. The dotted line indicates the median of ancestral strain, and the numbers beneath the line names indicate the number of cells imaged. p-values indicate the results of a t-test when each line is compared to the ancestor. **** p ≤ .0001, *** p ≤ 0.001, ** p ≤ 0.01, * p ≤ 0.05, ns = not significant. **B**. Distributions of cell volume with filamentous cells removed (cells with a volume larger than 3x the median for that line). **C**. Increase in volume is more strongly correlated with cell length compared to cell width. Each dot represents one cell. **D**. Relationship between the median volume with filaments removed and the total number of molecules of RNA per CFU. E. Correlation between total RNA per CFU for each replicate of each line. F. Correlation between the median cell volumes as determined in this work and cell volumes determined in Grant et al. 2020, figure 5. Error bars indicate the 25^th^ and 75^th^ quantiles of our data.

**Figure S3:**
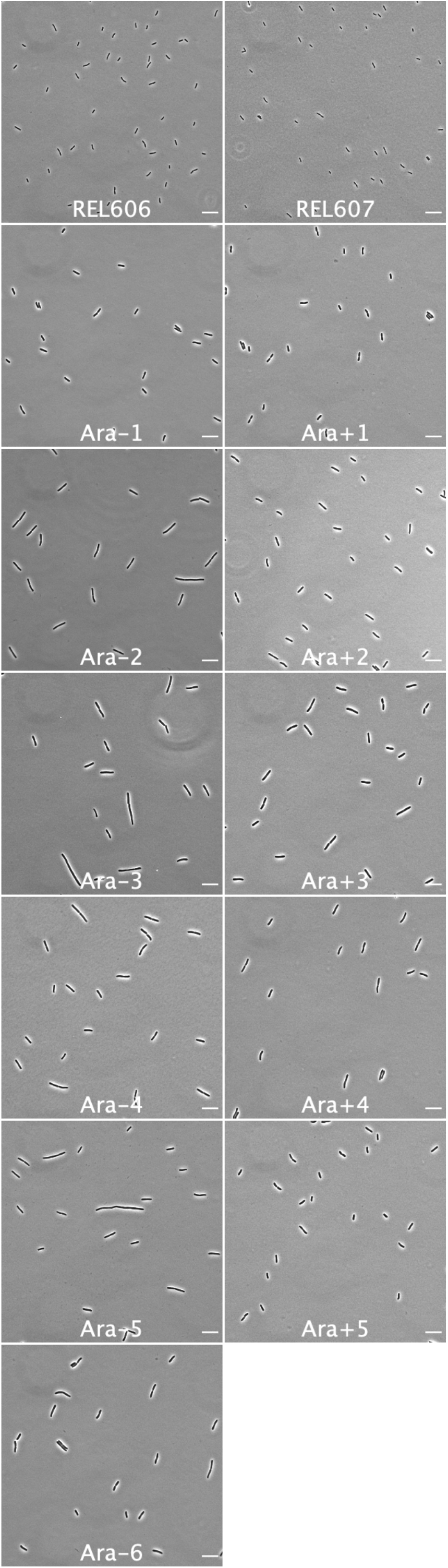
Representative phase contrast images of each of the lines used in this study. Scale bar is 3um.

**Figure S4:**
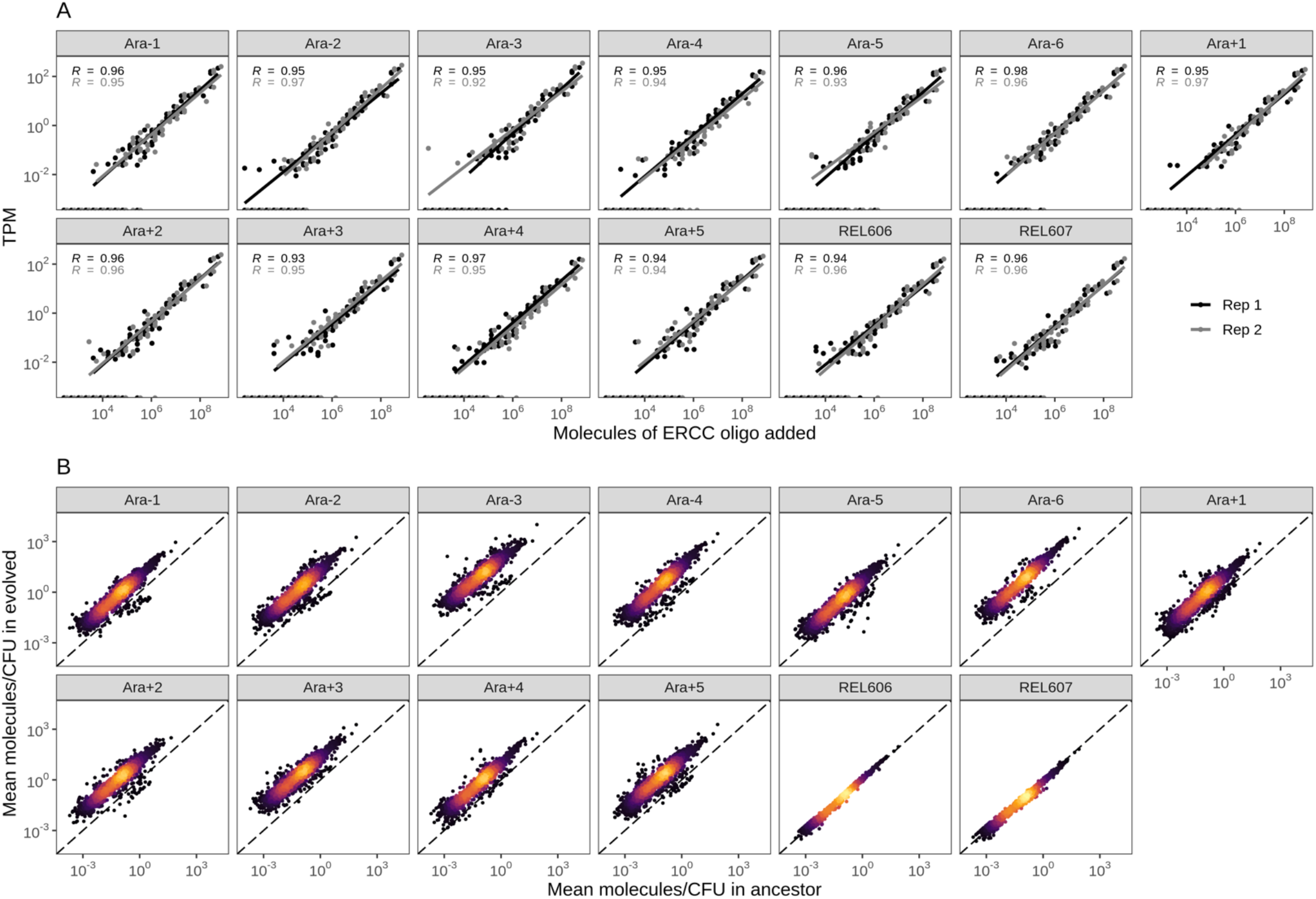
**A**, linear models relating the number of molecules of each ERCC control sequence added to their RNA-seq TPM (transcripts per million) in each line. **B**, changes in the absolute number of mRNAs/CFU in each line relative to the ancestor. The values plotted are the average between 2 replicates of the evolved line and both replicates from both ancestors (4 in total). **A.** Spike-in RNA control abundances are correlated with their estimates in sequencing data. Linear models relating the number of molecules of each ERCC control sequence added to their RNA-seq TPM (transcripts per million) in all RNA-seq samples. **D**. Most genes have a higher absolute expression in evolved lines. Changes in the absolute number of mRNA molecules per CFU (colony forming unit) in the 50,000th generation of each line relative to the ancestor. The values plotted are the average between 2 replicates of the evolved lines and both replicates from both ancestors. REL606 and REL607 are ancestral strains.

**Figure S5:**
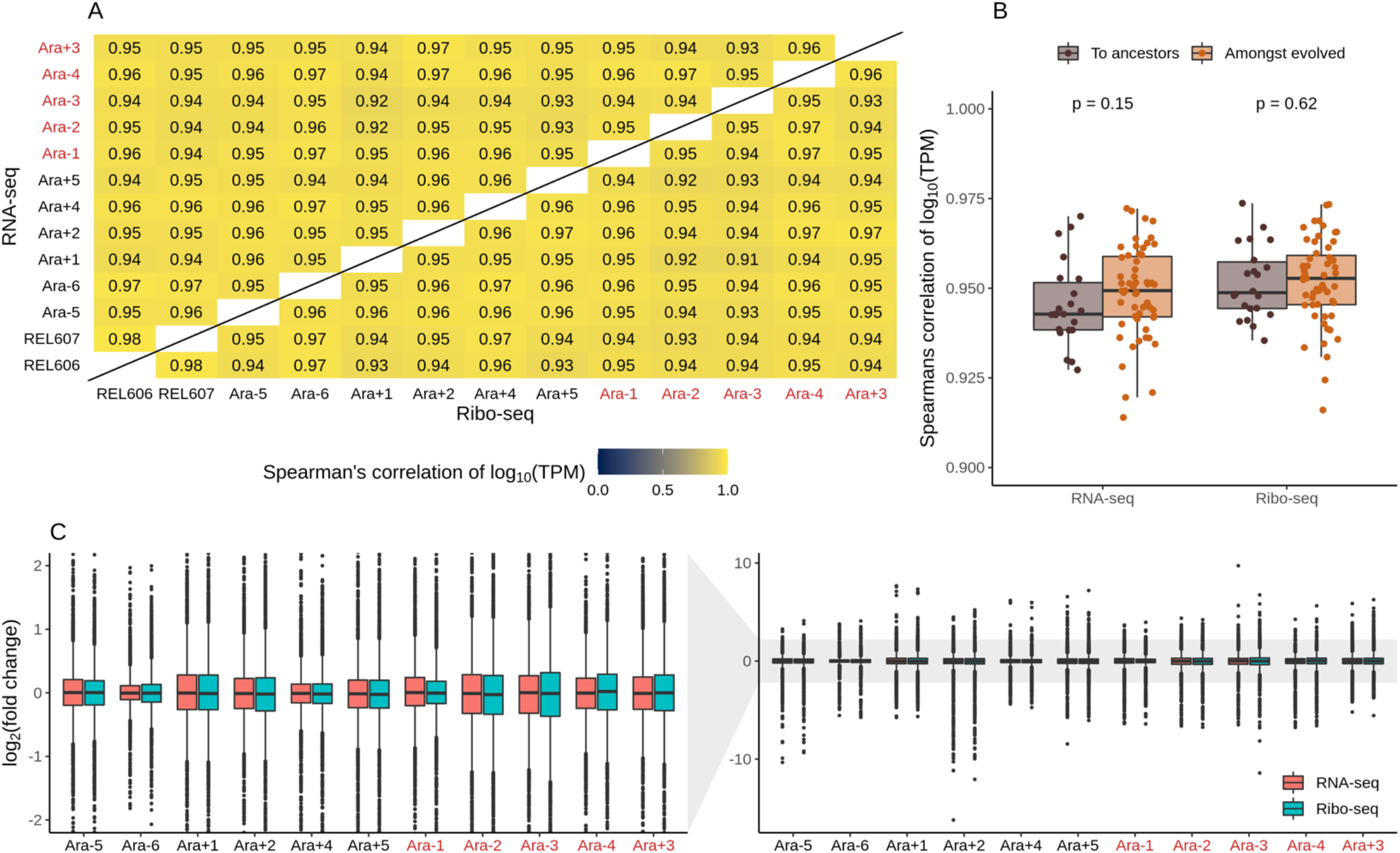
**A**. Pairwise correlations between expression levels of genes across lines based on log_10_(TPM). The upper triangle shows RNA-seq data, and the lower triangle indicates Ribo-seq data. **B**. Distributions of pairwise correlations between evolved lines and ancestors (purple) and amongst evolved lines (orange). **C**. Distributions of all DESeq2 fold-changes for both sequencing methods for all lines. The left panel is a zoom of the right panel.

**Figure S6:**
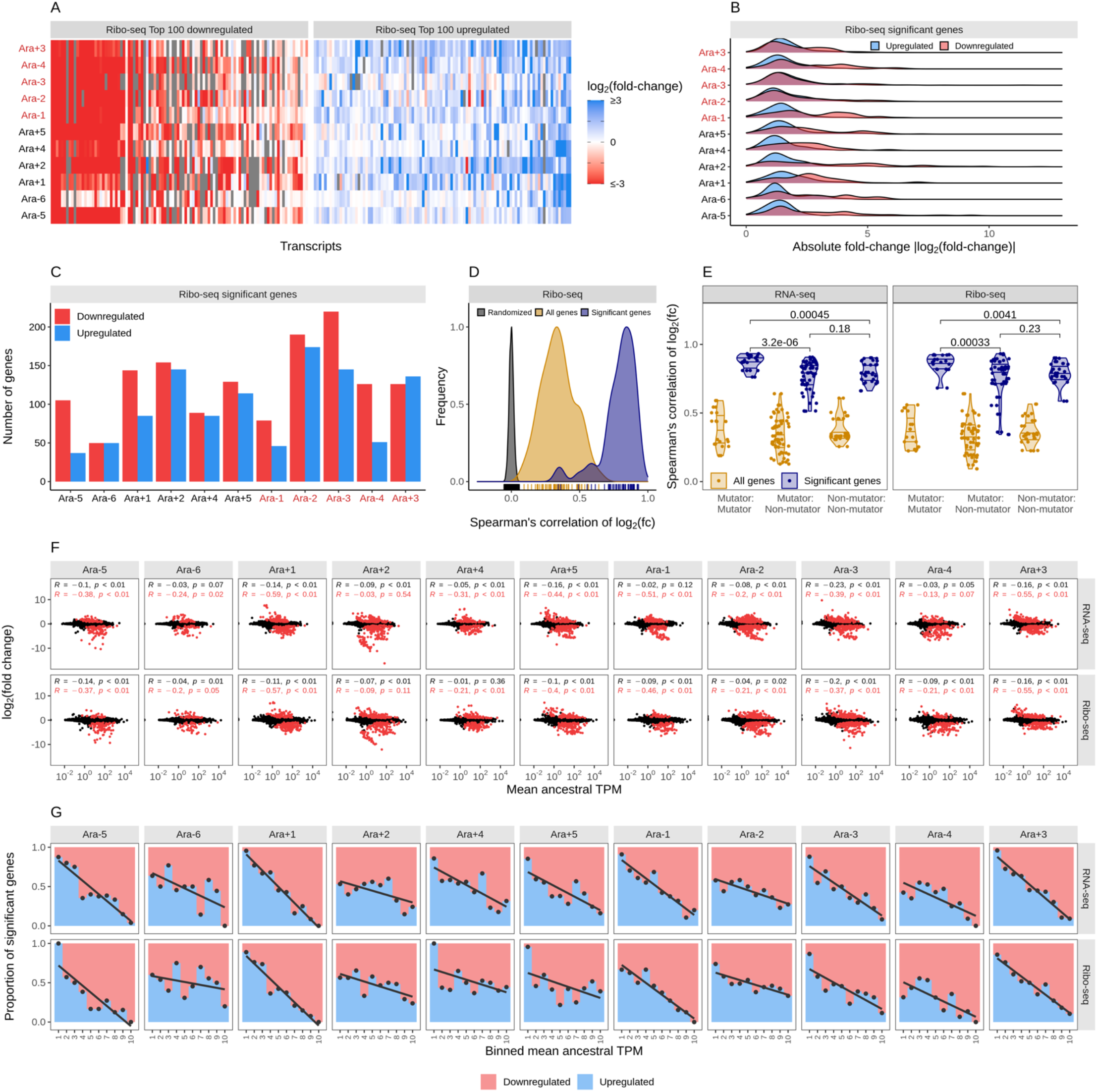
**A**. Parallelism in expression changes across evolved lines. The fold-changes of top 100 down and upregulated genes in each of the lines in the Ribo-seq datasets. Genes are ordered from left to right in order of increasing mean fold-change across evolved lines. Gray bars represent gene deletions. **B**. Downregulated genes have larger effect sizes than upregulated genes. Distribution of statistically significant fold-changes in Ribo-seq data in each line. Statistical significance was based on DESeq2 results using q ≤ 0.01. **C**. The number of significantly down and upregulated genes in each line. **D**. Pairwise correlations of evolved lines based on all (yellow curve) or statistically significant (blue curve) Ribo-seq fold-changes. Each of these curves is significantly different from a distribution based on correlations made after randomizing the fold-changes (grey curve) within each line (p ≤ 0.01, t-test). **E.** Pairwise-correlations between fold-changes in expression levels of genes based on their mutator status. **F**. Fold-changes in expression levels of genes in evolved lines scale negatively with their ancestral expression levels. The relationship between ancestral TPM in both RNA- and Ribo-seq datasets corresponding fold-changes across all lines. The black dots represent all the points (all genes), and the red dots represent significantly altered genes. **G.** Genes with high ancestral expression are typically downregulated. The panel shows the proportion of differentially expressed genes that are up/down-regulated as a function of ancestral expression (TPM).

**Figure S7:**
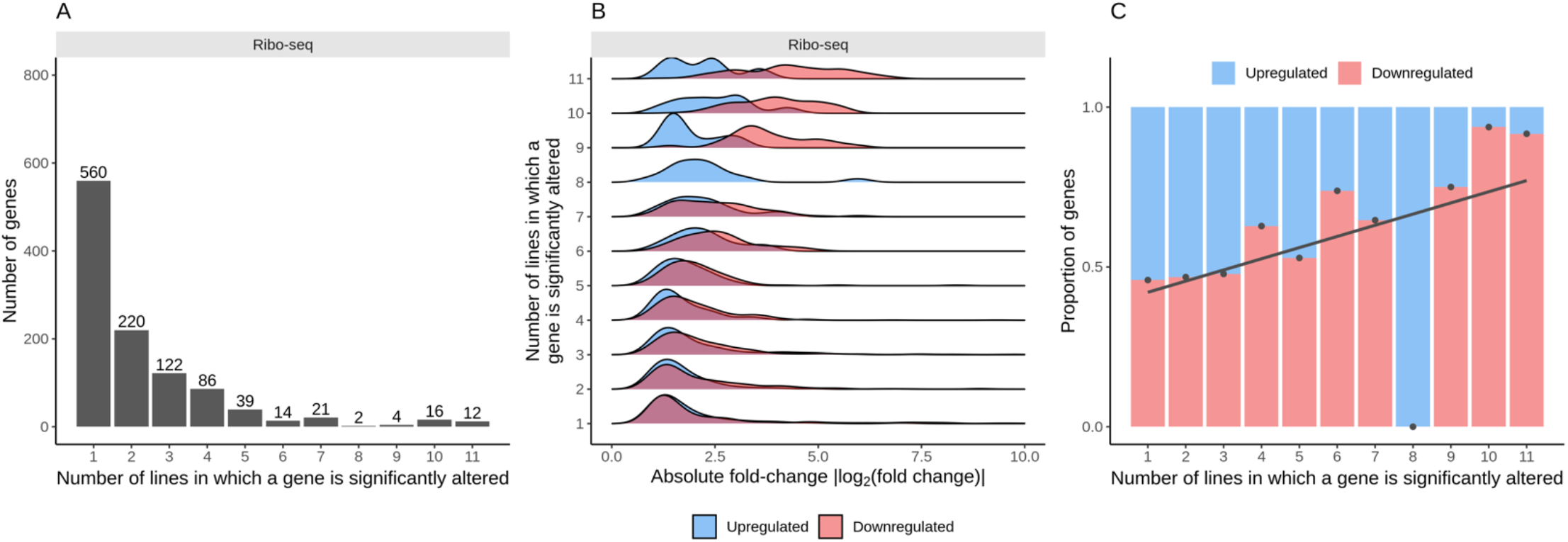
**A**. The number of evolved lines in which a gene’s expression level was significantly altered (q ≤ 0.01) was based on the DESeq2 results for the Ribo-seq dataset. **B**. Frequency downregulated genes have larger effect sizes than upregulated genes. Distributions of the Ribo-seq fold-changes for the genes. **C**. Frequently altered genes are typically downregulated. The proportion of up and downregulation of genes in the Ribo-seq dataset as a function of their frequency of expression changes across lines.

**Figure S8:**
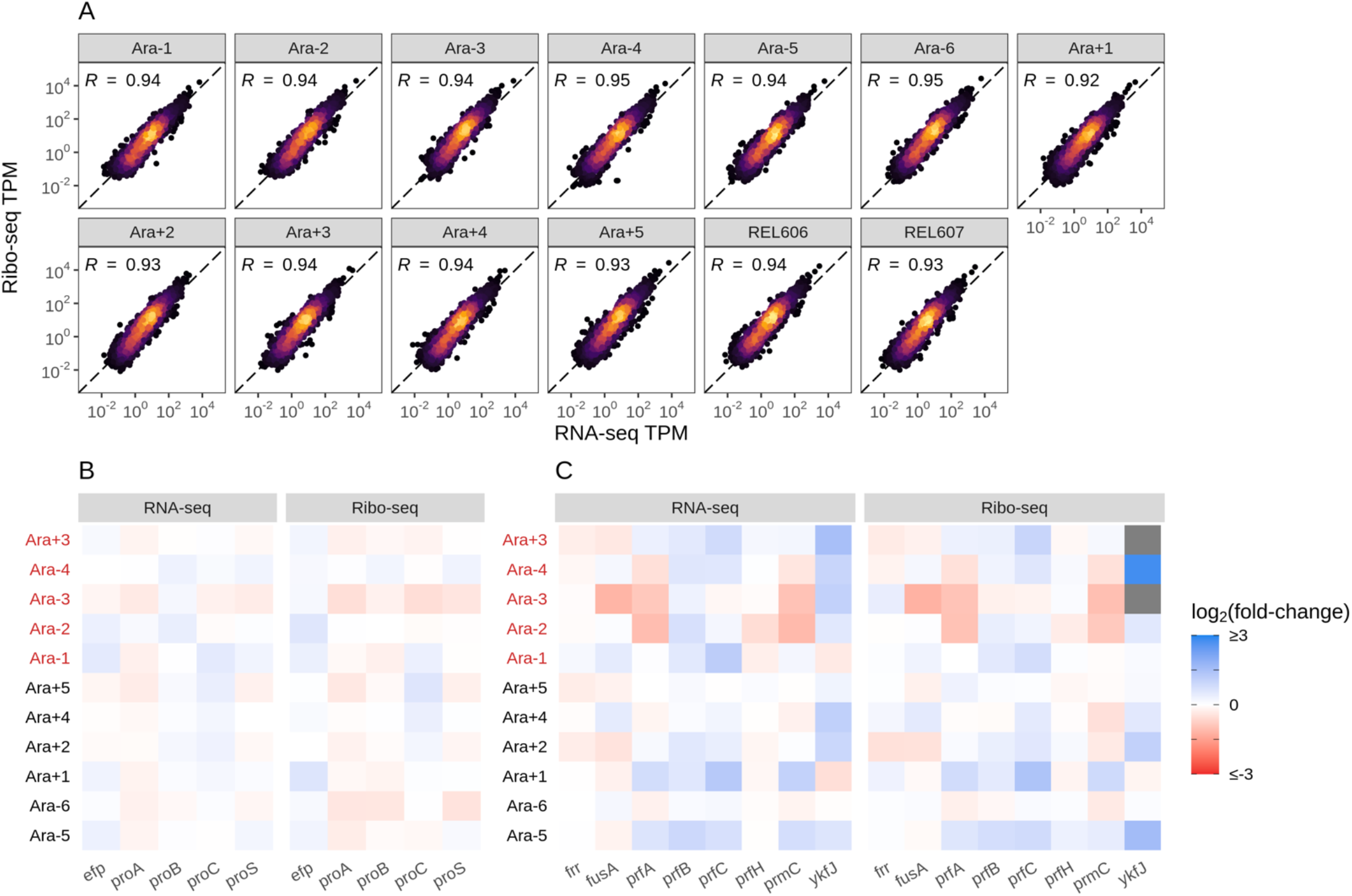
**A**. Translational changes are positively correlated with transcriptional changes. The relationship between RNA-seq and Ribo-seq TPM across all evolved lines. The TPMs are averaged between the replicates. **B**. Fold-changes in expression levels of genes involved in proline biosynthesis. **C**. Fold-changes in expression levels of translation termination factors and related genes.

**Figure S9:**
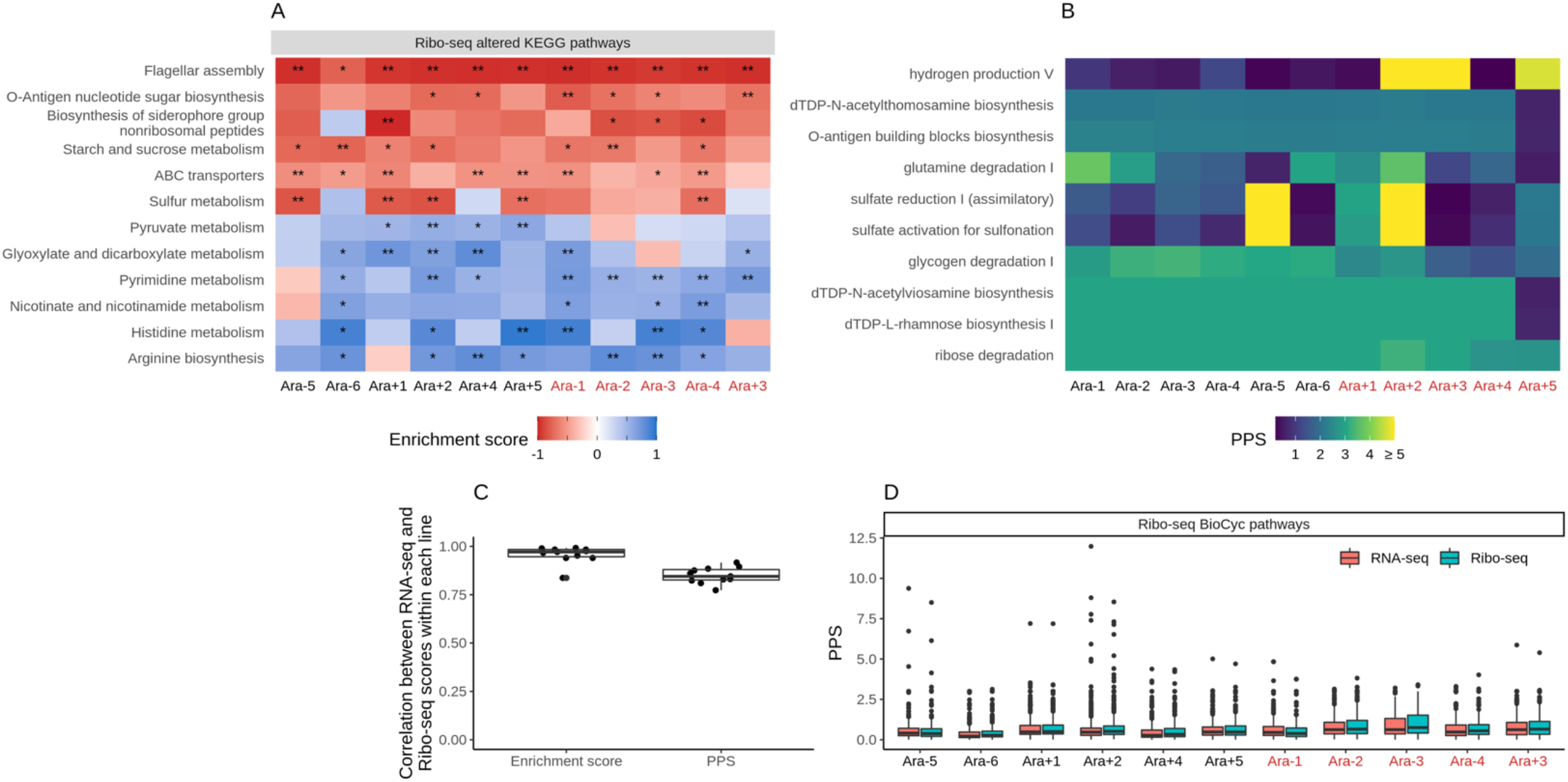
**A.** Parallel changes in functional categories. KEGG enrichment scores from the Ribo-seq data. Enrichment score represents the degree to which a pathway was up (positive) or downregulated (negative). Functional categories are ordered by increasing mean enrichment score across the lines. Enrichment score represents the degree to which a pathway was up (positive) or downregulated (negative). **B**. Pathway perturbation score (PPS) calculated from Ribo-seq fold changes. Higher PPS indicates larger degrees of alteration but does not indicate directionality. **C**. Pairwise correlations of KEGG enrichment scores for all pathways that were significantly altered in at least one line. **D**. Distribution of PPS scores in both RNA-seq and Ribo-seq datasets across all lines.

**Figure S10:**
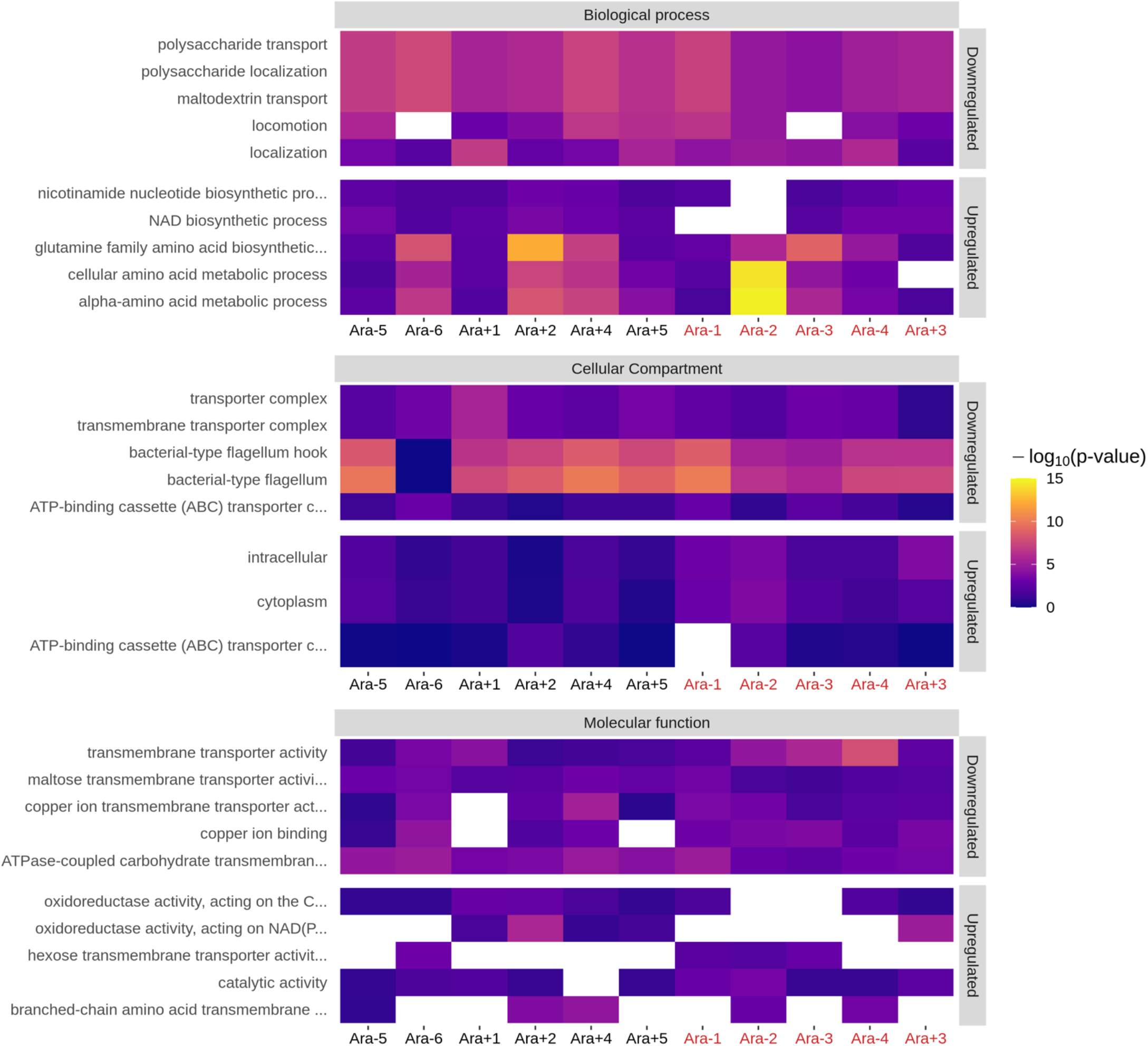
The top 5 up and downregulated GO categories for each ontology term. For each ontology, only terms with a p-value ≤ 0.01 based on Fisher’s exact test in at least 4 lines were considered. White spaces indicate that a particular category was not significantly altered in a line.

**Figure S11:**
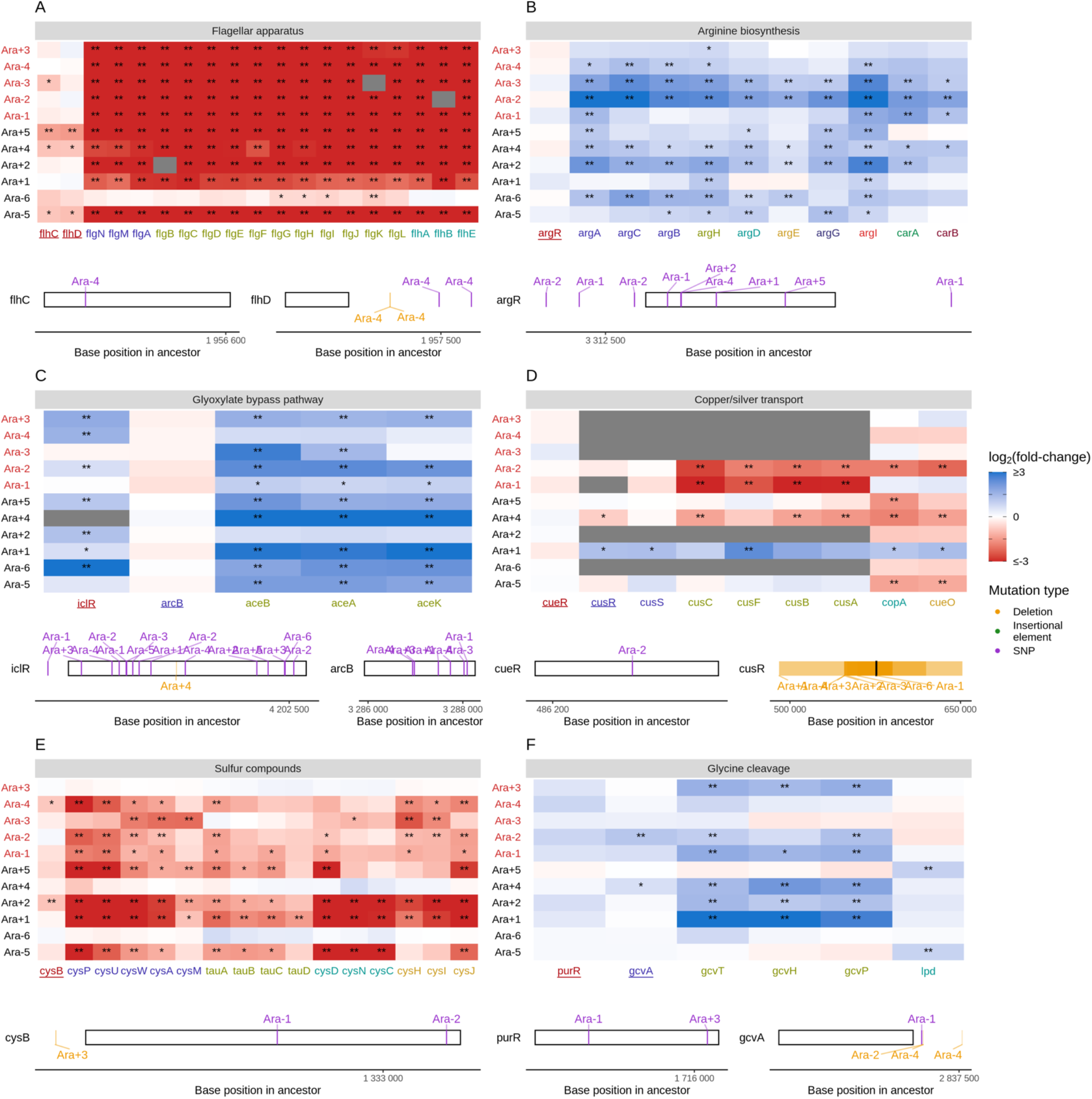
**A-F**. Mutations in transcriptional regulators lead to parallel changes in gene expression (RNA-seq). Gene names in each category are colored based on their operon membership. Transcription factors for each class of genes are underlined. Asterisks indicate statistical significance of fold-changes, ** q ≤ 0.01, * q ≤ 0.05. Grey panels in the heatmap indicate gene deletion. Lower panels show the type and location of mutations in each transcription factor.

